# Coupling of Nuclear Translocation to Cell Size Promotes Robustness to Fluctuations in YAP/TAZ Concentration

**DOI:** 10.1101/2023.02.06.527281

**Authors:** Ian Jones, Mar Arias-Garcia, Patricia Pascual-Vargas, Melina Beykou, Lucas Dent, Tara Pal Chaudhuri, Theodoros Roumeliotis, Jyoti Choudhary, Julia Sero, Chris Bakal

## Abstract

The concentration of many transcription factors exhibit high cell-to-cell variability due to differences in synthesis, degradation, and cell size. How these factors are robust to fluctuations in concentration is poorly understood. Here we quantified the single cell levels of the YAP/TAZ transcriptional co-activators in parallel with cell morphology for over 400,000 single cells across 17 cell lines. We show the whole cell concentration of YAP/TAZ sub-scales with respect to size as cells grow during proliferation. However, the mean nuclear concentration of YAP/TAZ remains constant during the cell cycle. Theoretical modelling demonstrates that the extent to which whole cell YAP/TAZ dilutes in single cells during proliferative growth dictates the variability of YAP/TAZ levels across the population. Integrative analysis of imaging and proteomic data show the average nuclear YAP/TAZ concentration is predicted by differences in RAS/MAPK signalling, focal adhesion maturation, and nuclear transport processes. We developed a statistical framework capable of discriminating between perturbations that affect YAP/TAZ directly, or via changes in morphology. Deployment of these models on genetic screening data or small-molecule treatments reveal that inhibition of MEK, CDK4/6, LATS and RhoGTPases decouple nuclear YAP/TAZ from cell morphology by regulating nuclear translocation. Thus signalling activity couples size changes to YAP/TAZ translocation; leading to a stable pool of nuclear YAP/TAZ during proliferation.

**Significance Statement:** Many proteins dilute/concentrate with changes in cell size. It is unclear how robustness in cell signalling emerges across differently sized cells, with varying intracellular protein concentrations, over generations. Here, we have shown that despite whole cell dilution of the transcriptional co activators YAP/TAZ with increasing size, a steady-state nuclear concentration distribution is maintained across the population. Thus nuclear transport promotes robustness of signal response in the face of a dwindling cytoplasmic YAP/TAZ levels. An integrative approach revealed that focal adhesions, RAS/MAPK and nuclear import contributes to the the maintenance of YAP/TAZ nuclear levels. Cells appear to have evolved systems to ensure robustness against alterations to cell size during the cell cycle.

---

Signalling networks couple transcriptional regulation to the integrated detection of environmental cues. A common ‘motif’ in these networks is the sequestration of transcription factors by inhibitory complexes. In the presence of an environmental cue, transcription factors are released from these complexes and interact with DNA to engage specific programmes (1–3). When TFs and inhibitors are at sub-saturating levels, this allows transcription to be tightly coupled to environmental flux. For example, dilution of the inhibitory protein RB1 as cells grow is one mechanism by which E2F activity is coupled to size (4, 5). In animal cells especially, inhibitory sequestration of transcription factors often occurs in the cytoplasm; and release from inhibition allows TFs to translocate into the nucleus. For example, the active degradation of inhibitors such as APC or IKB in the cytoplasm allows the translocation of transcription factors such as Beta-Catenin or NFKB into the nucleus (6, 7); coupling cues such as adhesion and stress to transcription respectively. The concentration of both inhibitors and transcription factors in the cytoplasm thus informs the response of transcription factors to upstream signals (4).

It is now clear that the concentration of many cellular molecules varies between cells, even within an isogenic population (8). Such variability can be due to both extrinsic and intrinsic stochastic differences in protein synthesis, but also due to differences in cell size and shape (8–11). However, cell populations and tissues exhibit largely robust and predictable behaviour despite such fluctuations; i.e. are largely uniform in the output of their signalling activity. Perhaps the best example of which is the control of size uniformity during proliferation, such as during organ and tissue morphogenesis (12–14). But how cells are robust to fluctuations in protein concentration is poorly understood.

In the context of a sub-scaling protein, one means to provide robustness would be to couple a synthesis step to cell growth, such that a surge in synthesis offsets the effects of dilution. Indeed, such a system is believed to underpin the maintenance of RB1 (yeast WHI5) concentration across consecutive cycles (4). In the Rb1/Whi5 context, this coupling is complemented by saturating DNA with RB1/WHI5 (and degrading the excess) prior to division (15); where the amount of protein inherited by either daughter is proportional to the DNA content - and thus size - of the cell. Indeed, a similar system applies to the partitioning of KRP4 in Arabidopsis (16). Importantly, these systems can do nothing to constrain the effects of dilution within a cell cycle. However, it is unclear how the effects of dilution may be mitigated in other biological contexts.

YAP and TAZ (henceforth YAP/TAZ) are conserved key regulators of animal cell growth and proliferation. As transcriptional co-activators, YAP/TAZ are inhibited by sequestration in the cytoplasm, where nuclear translocation (activation) is triggered by soluble, mechanical, and geometrical cues (17–25). For example, changes in cell shape are coupled to the signalling dynamics of Rho GTP Exchange Factors (RhoGEFs), Rho GTPase Activating Proteins (RhoGAPs) and their downstream effector Rho GTPases. RhoGEFs, RhoGAPs, and Rho GTPases regulate YAP/TAZ translocation both by regulating YAP/TAZ signalling directly and by affecting cell shape/size (indirectly) (18, 22, 26–32). The coupling of YAP/TAZ dynamics to cell shape provides a mechanism that allows cells to position themselves during development, or to sense and respond to disruption in tissue structures. (26, 33– 35). YAP/TAZ are also regulated by the Hippo pathway, whereby the LATS (LATS1 and LATS2) kinases phosphorylate YAP/TAZ, preventing nuclear translocation by enabling 14-3-3 binding.

Through quantitative analysis of 100,000’s of single cells, from 17 cell lines, we have demonstrated that the whole cell and cytoplasmic YAP/TAZ concentration sub-scales with cell size in G1 and G2. YAP/TAZ synthesis was dramatically upregulated near S-Phase in a size and ploidy dependent manner. Crucially, we observed that the nuclear YAP/TAZ concentration distribution was constant across the population when binned by cell size, implying continual nuclear import in the background of depleting whole cell YAP/TAZ concentration. Through integrative analysis of proteomic data, we found that YAP/TAZ nuclear transport is predicted by the phosphorylation state of RAS/MAPK, focal adhesion, and nuclear transport components; suggesting a role for these systems in coupling cell/cytoplasm size to nuclear import. Using a novel statistical framework, we show that RAS/MAPK, CDK4/6, and RHOA affect YAP/TAZ translocation directly. Whereas inhibition of genes such as LATS1 and LATS2 affect both size and translocation. Taken together, our work defines a system to ensure the robustness of cell signalling to changes in protein concentration.

## 1. Results

### A. YAP/TAZ concentration decreases with increasing cell size

We quantified single cell size and the concentration/abundance distributions of F-actin and cytoplasmic/nuclear YAP/TAZ, first in 30,000 single cells from nine breast cell lines (Table 1) (Fig. 1A). While ostensibly YAP specific, the antibody we used partially recognises TAZ though to a much lesser extent (23, 36). We initially investigated whether average YAP/TAZ intensities varied with cell size across our panel but found no evidence of a correlation with the nuclear or whole cell signal (Fig. 1B/C). However, there was a clear linear relationship between the whole cell and nuclear YAP/TAZ mean intensities (Fig. 1D). Thus high expression of YAP/TAZ correlates with more nuclear import. This relationship was also observed in single cells within each cell line (Fig. 1E). Strikingly, when investigating whether area predicts whole cell YAP/TAZ in single cells, we observed a clear negative correlation. Meaning that whole cell YAP/TAZ dilutes/degrades as the cells grow (Fig1F). This prompted us to more formally investigate the decrease of whole cell YAP/TAZ within each cell line.

**Table 1.** Cell Line Information: Gene cluster: Lu = luminal, BaA = Basal A, BaB = Basal B. ER/PR/HER2: +/ from protein and mRNA expression; [] inferred from mRNA expression; M = mutant, WT = wild-type. MDA 231-LM2-4175* cells are the highly metastatic subpopulation 4175 from MDA-MB-231 (23, 37–39).

| Cell Line | Genetic Subtype | ER | PR | Her2 |
| --- | --- | --- | --- | --- |
| MCF7 | Lu | + | [+] | – |
| T47D | Lu | + | [+] | – |
| BT474 | Lu | + | – | + |
| SKBR3 | Lu | – | [–] | + |
| HCC1954 | BaA | – | [–] | + |
| MDA-MB-468 | BaA | [–] | [–] | – |
| hs578T | BaB | – | [–] | – |
| MDA-MB-157 | BaB | – | [–] | – |
| MDA-MB-231 | BaB | – | [–] | – |
| SUM149 | BaB | [–] | [–] | – |
| SUM159 | BaB | [–] | [–] | – |
| MCF10A | BaB | – | [–] | – |
| JIMT1 | Unclassified | – | [–] | + |
| MDA-MB-231-LM2-4175* |  |  |  |  |

**Table 2.**
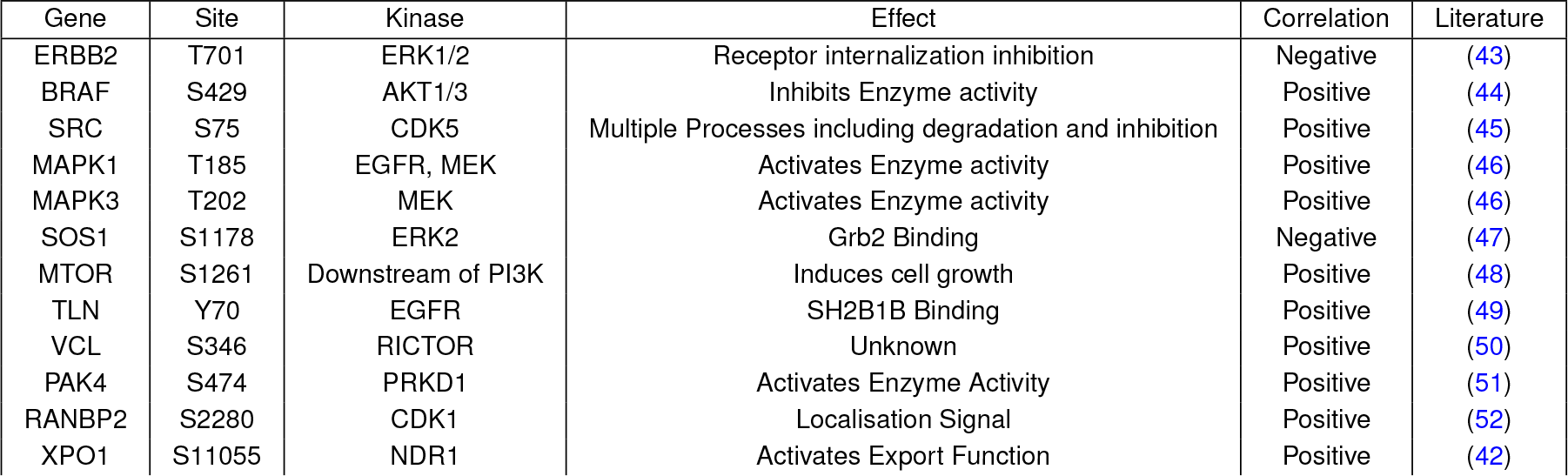
Select phosphorylation sites strongly predictive of the average nuclear YAP/TAZ concentration and/or Nuc/Cyto ratio across lines.

**Table 3.**
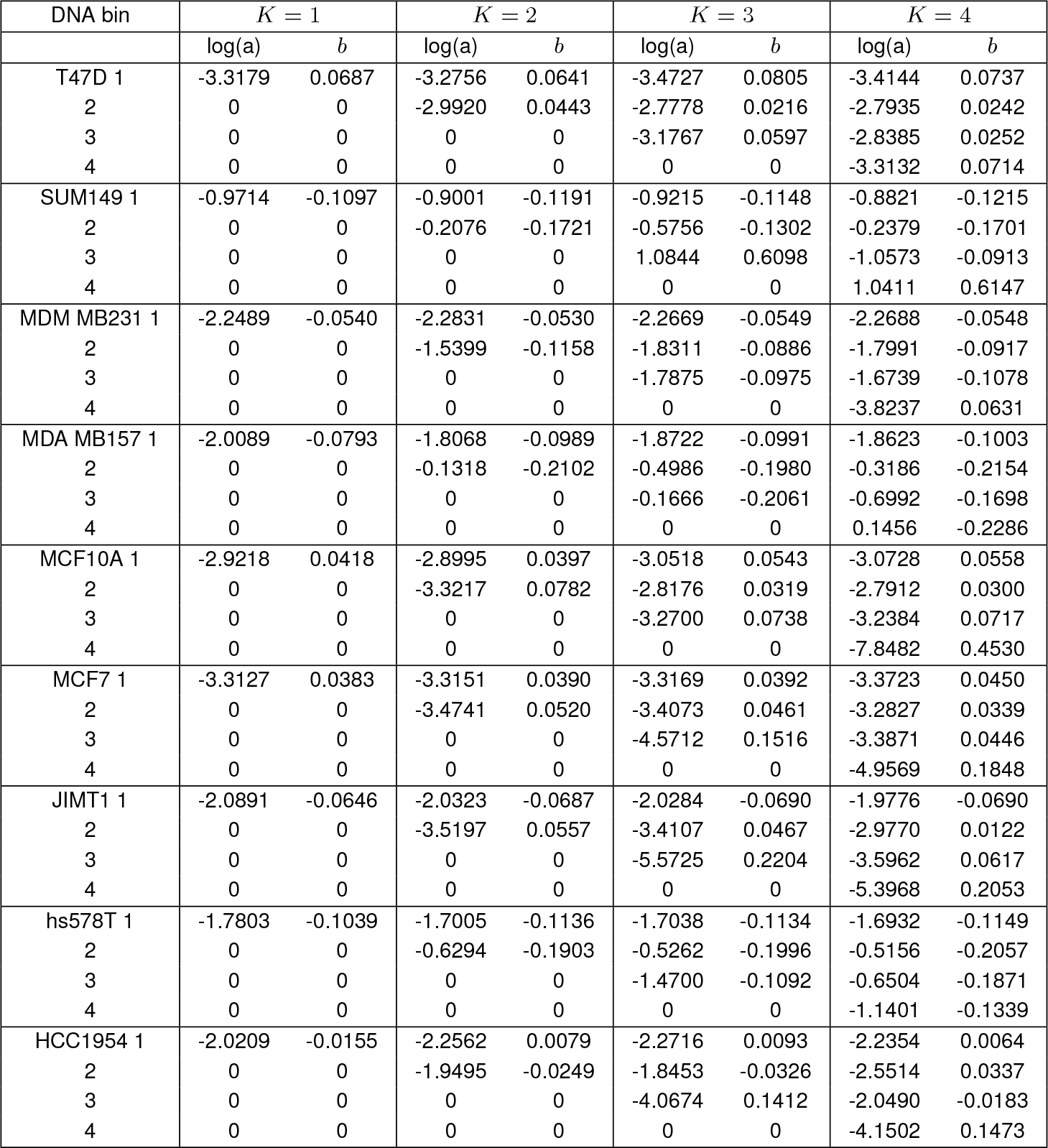
Actin Concentration Scaling: Scaling parameter values for each cell line across DNA bins within each ‘K’, the cluster number used in kmeans clustering on the integrated Hoechst intensity. Log(a) is proportional to the initial Actin concentration and b is the power to which Actin concentration scales with cell area.

**Table 4.**
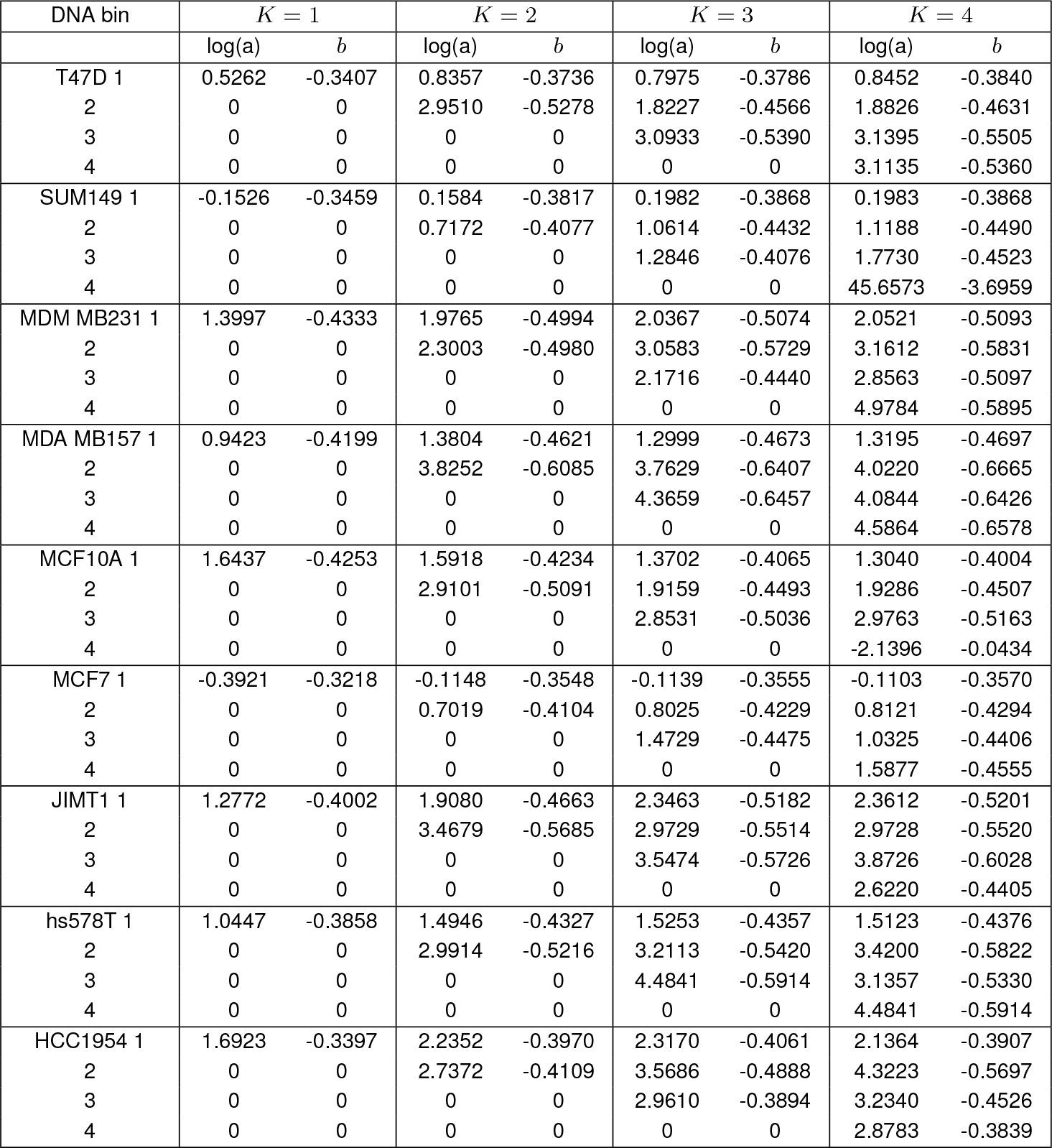
YAP/TAZ Concentration Scaling: Scaling parameter values for each cell line across DNA bins within each ‘K’, the cluster number used in kmeans clustering on the integrated Hoechst intensity. Log(a) is proportional to the initial YAP/TAZ concentration and b is the power to which YAP/TAZ concentration scales with cell area.

**Table 5.**
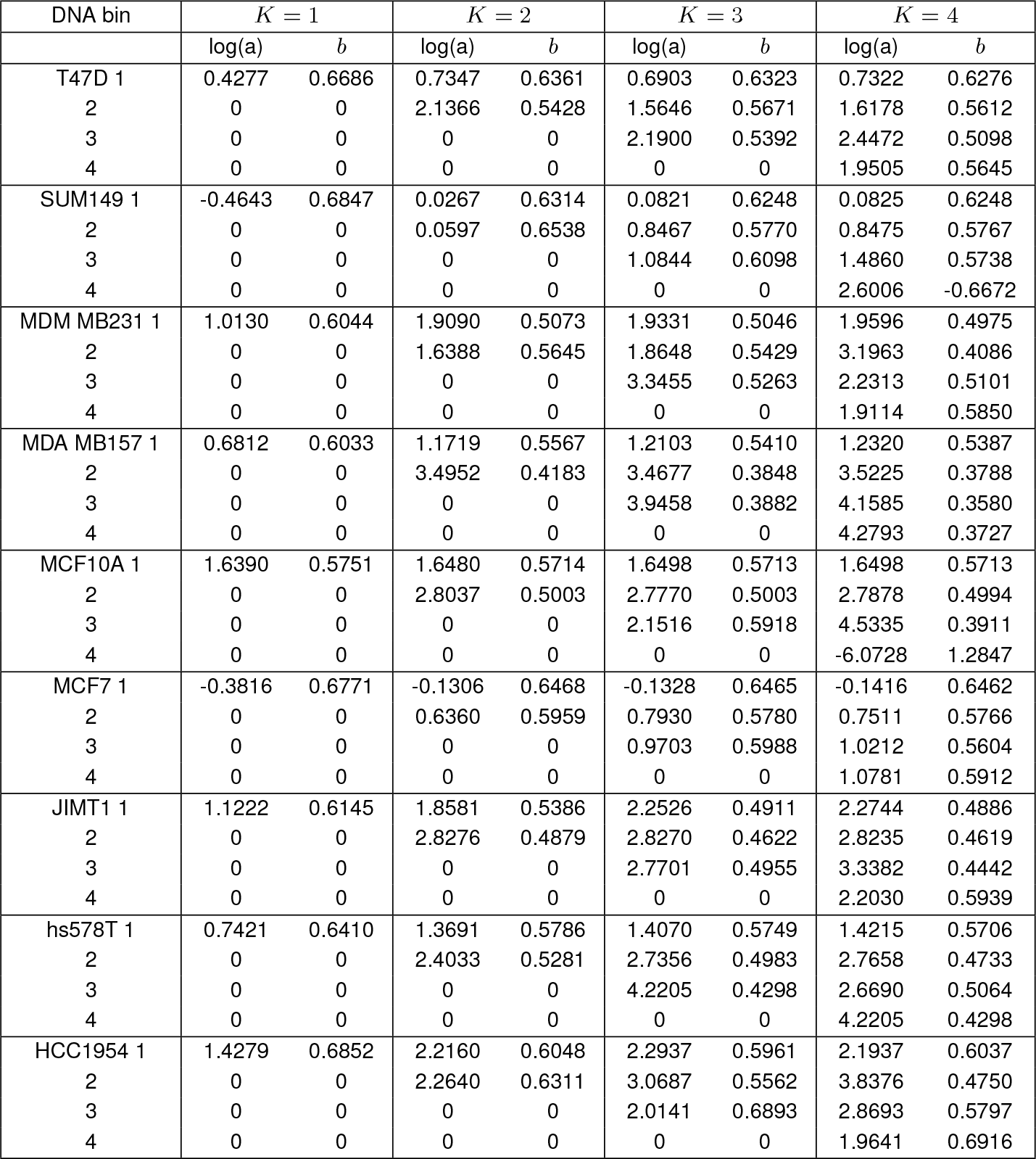
YAPTAZ Abundance Scaling: Scaling parameter values for each cell line across DNA bins within each ‘K’, the cluster number used in kmeans clustering on the integrated Hoechst intensity. Log(a) is proportional to the initial YAP/TAZ abundance and b is the power to which YAP/TAZ abundance scales with cell area.

**Table 6.**
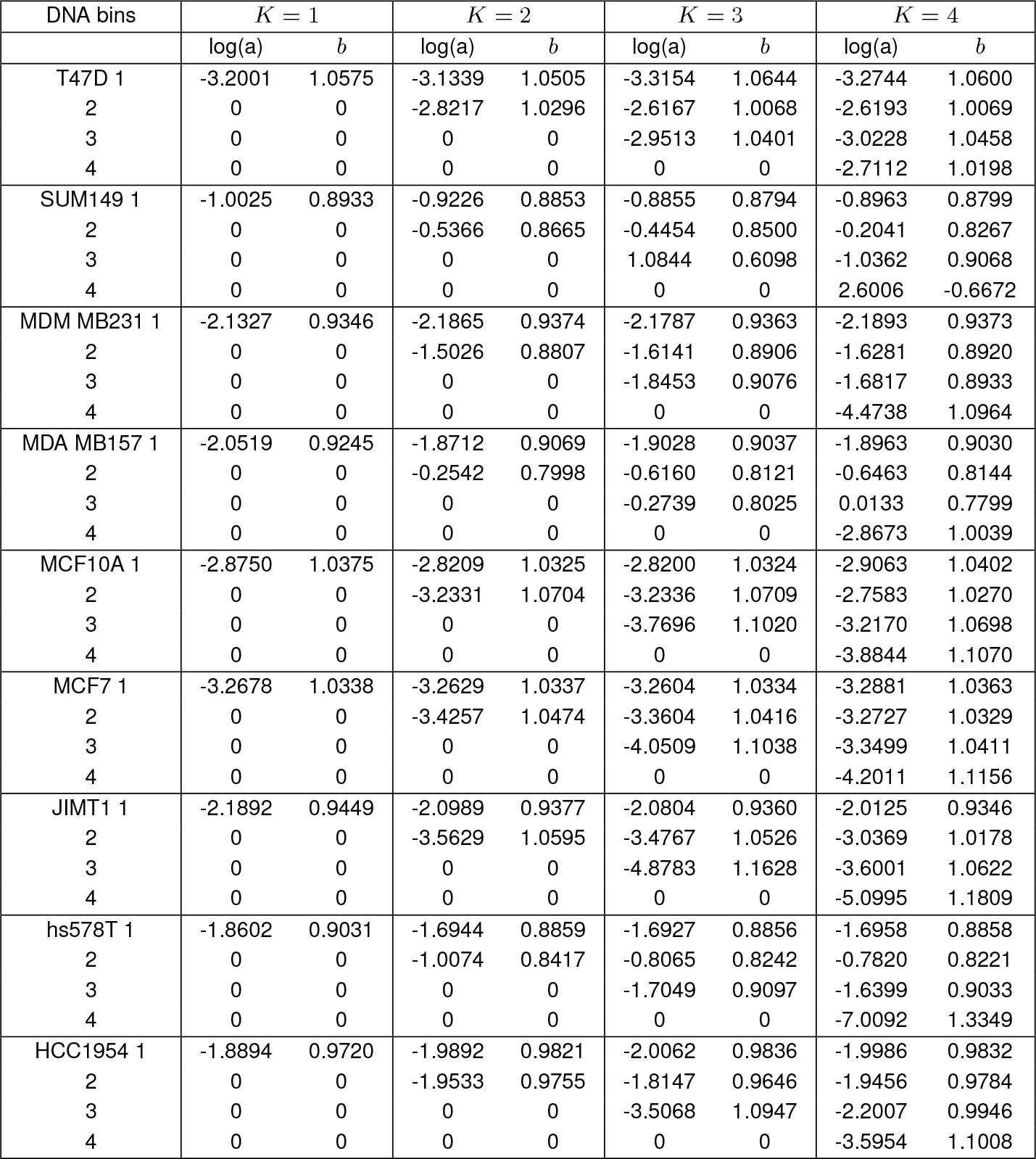
Actin Abundance Scaling: Scaling parameter values for each cell line across DNA bins within each ‘K’, the cluster number used in kmeans clustering on the integrated Hoechst intensity. Log(a) is proportional to the initial Actin abundance and b is the power to which Actin abundance scales with cell area.

**Table 7.**
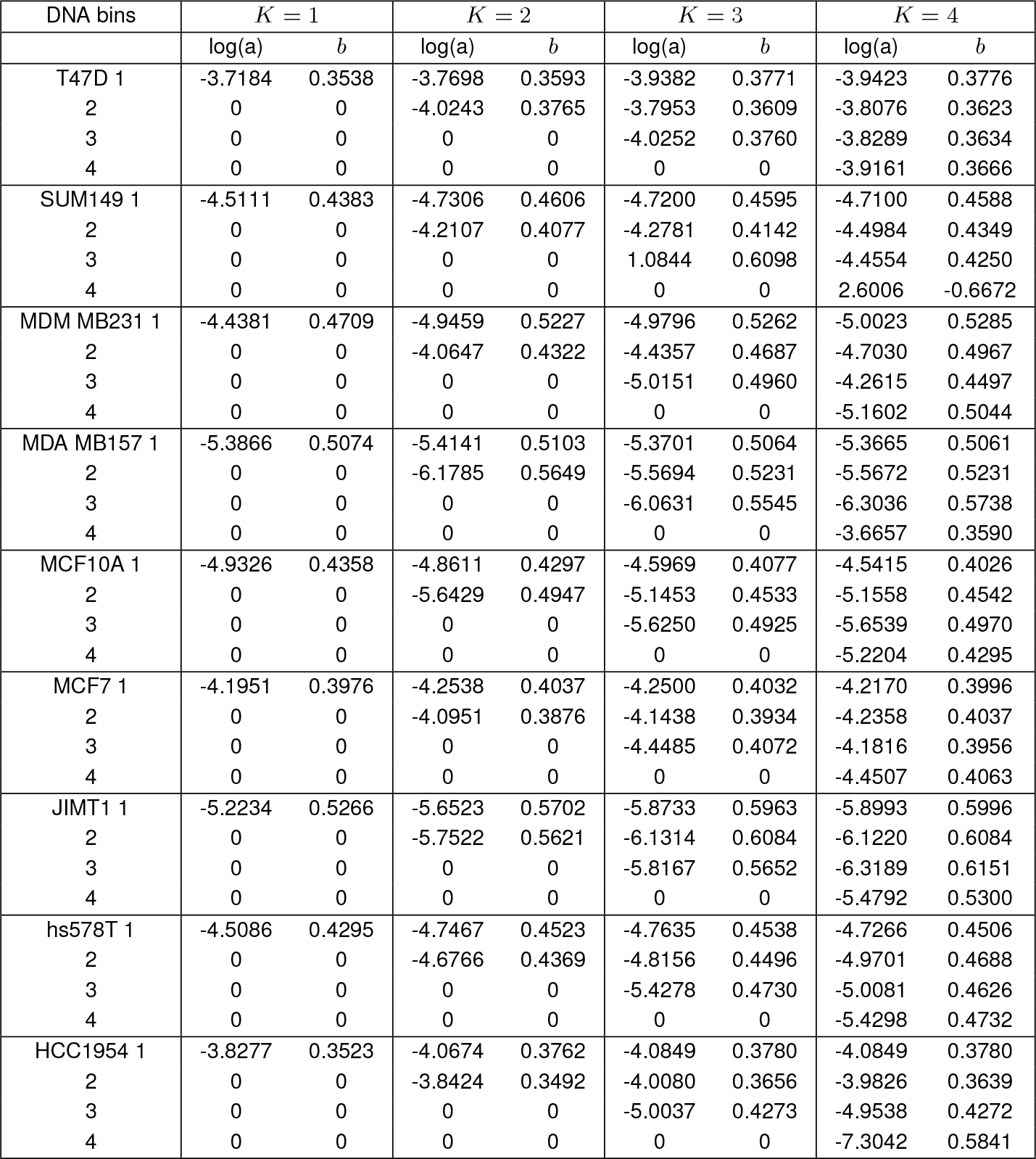
YAP/TAZ Ratio Scaling: Scaling parameter values for each cell line across DNA bins within each ‘K’, the cluster number used in kmeans clustering on the integrated Hoechst intensity. Log(a) is proportional to the initial YAP/TAZ ratio and b is the power to which YAP/TAZ ratio scales with cell area.

**Table 8.**
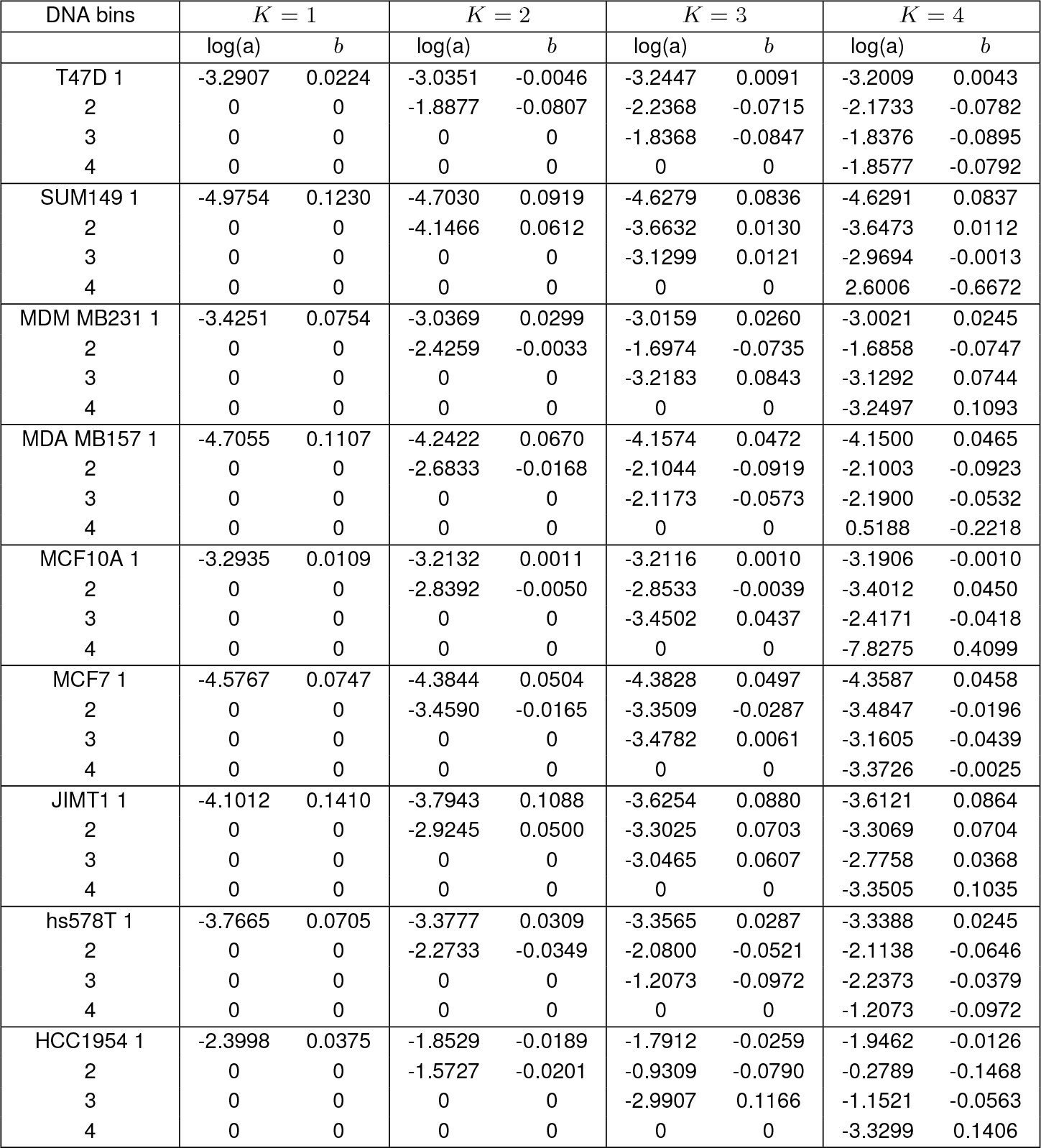
YAP/TAZ Nuclear Scaling: Scaling parameter values for each cell line across DNA bins within each ‘K’, the cluster number used in kmeans clustering on the integrated Hoechst intensity. Log(a) is proportional to the initial YAP/TAZ nuclear concentration and b is the power to which YAP/TAZ nuclear concentration scales with cell area.

**Table 9.**
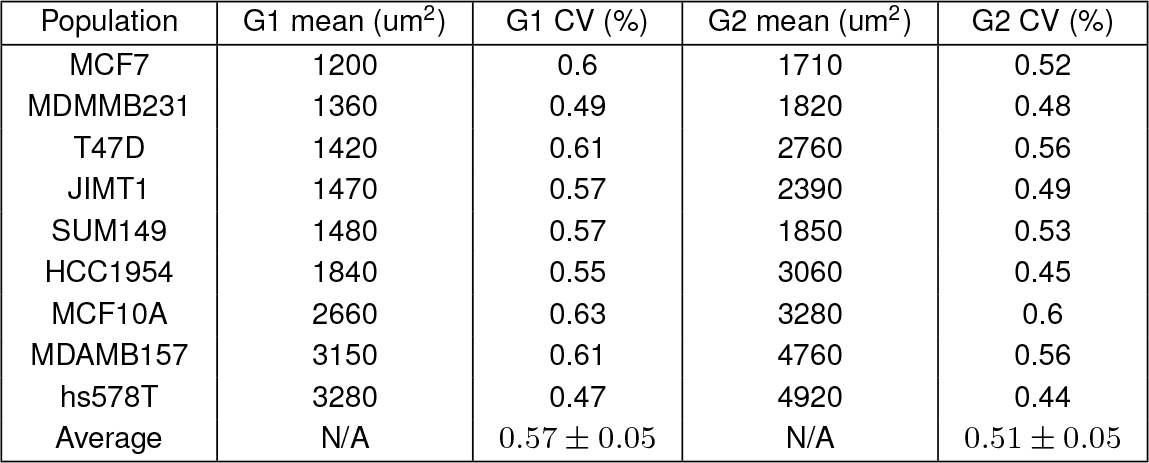
Population Characteristics: Cell area distribution statistics. ‘G1’ and ‘G2’ are defined via kmeans clustering on the integrated Hoechst intensity.

**Table 10.**
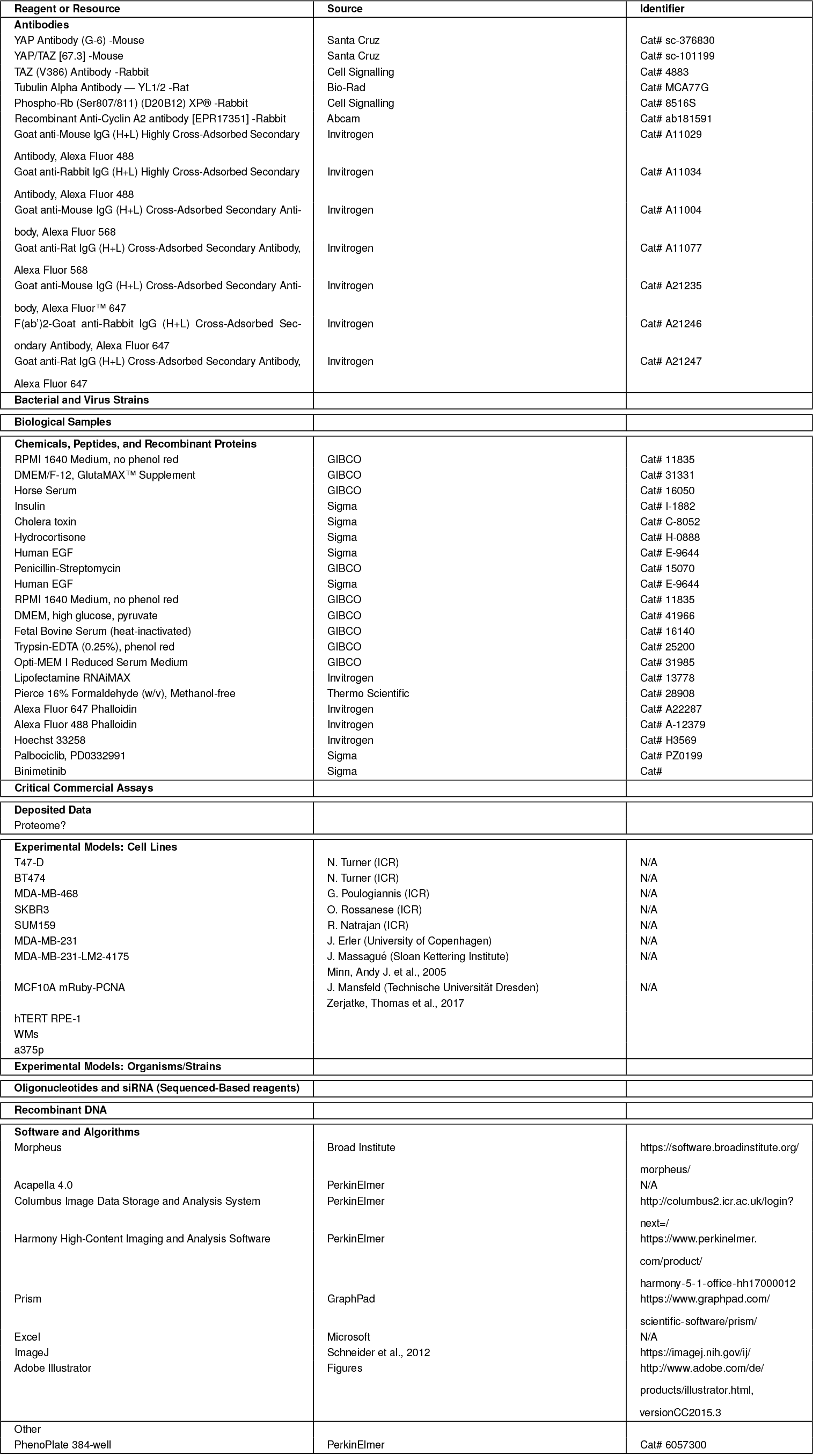
List of reagents and resources.

**Figure 1.**
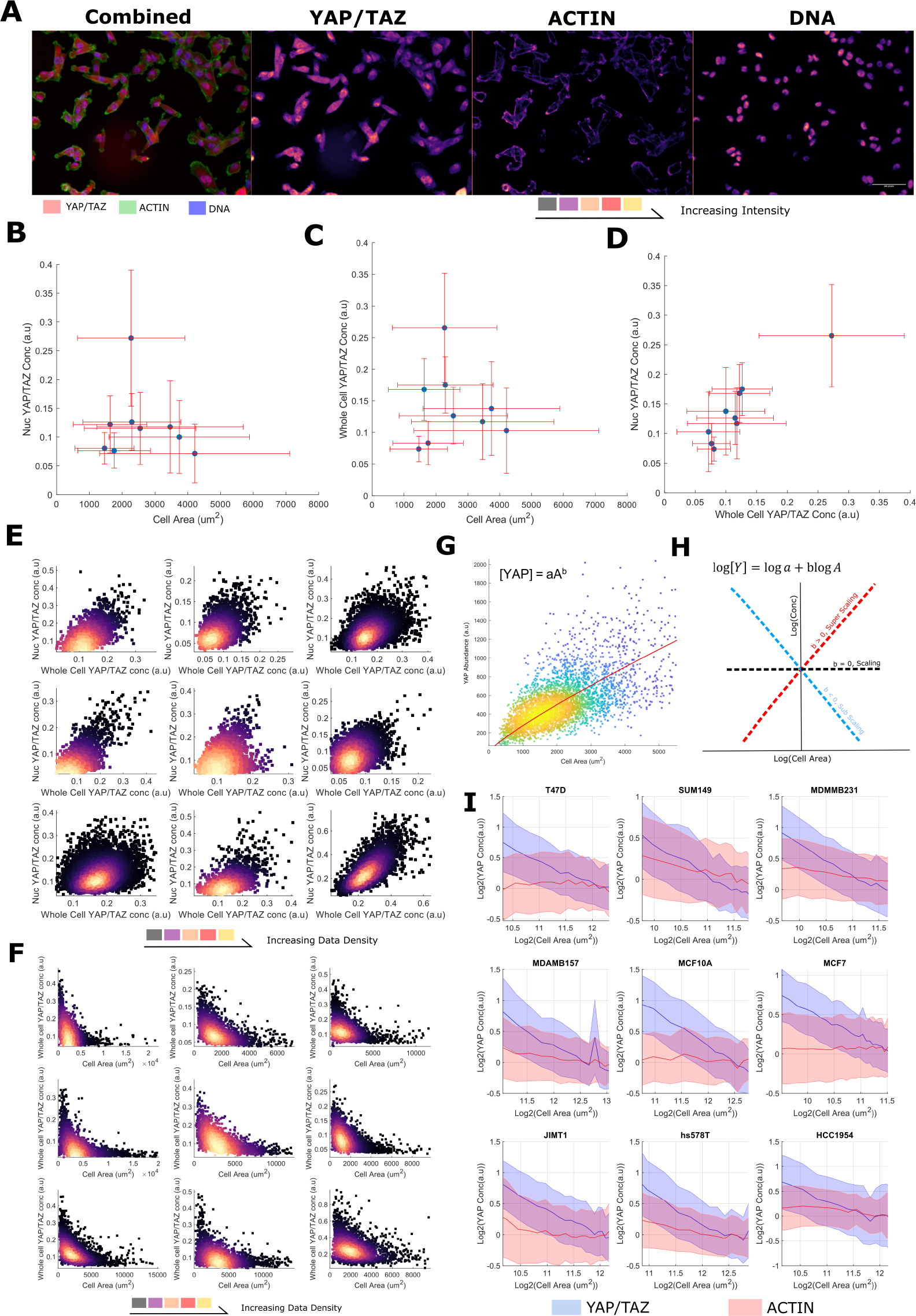
YAP/TAZ concentration decreases with increasing cell size A) (left) Representative image of MBA-MB-231 cells stained with YAP/TAZ (red), Phalloidin (an actin binding dye, green) and Hoescht (marking the DNA in blue). (Right) Individual colour channels seperated out from the original image. Colour is proportional to image intensity such that *black < purple < red < yellow*. The scale bar represents 50 um. B) The relationship between the nuclear YAP/TAZ concentration and cell area across cell lines. Error bars represent one standard deviation. C) The relationship between the whole cell YAP/TAZ concentration and cell area across cell lines. Error bars represent one standard deviation. D) The relationship between the whole cell YAP/TAZ concentration and the nuclear YAP/TAZ concentration across cell lines. Error bars represent one standard deviation. E) The relationship between the whole cell YAP/TAZ concentration and the nuclear YAP/TAZ concentration in single cells in each cell line. The colour is proportional to data density such that *black < purple < red < yellow*. F) The relationship between the whole cell YAP/TAZ concentration and cell area in single cells in each cell line. The colour is proportional to data density such that black ¡purple ¡red ¡yellow. G) YAP/TAZ ‘abundance’ (integrated intensity) as a function of cell area in T47D cells. We describe this relationship as a power law, y = aAb, shown in red. The colour of the data is proportional to the density of the data. H) Demonstration of how the log-log plot of size vs concentration is interpreted; a negative gradient corresponds to dilution with growth, positive indicates an increasing concentration and a flat relationship, perfect scaling with cell size. I) Log-log plots relating YAP/TAZ (blue) and Actin (red) concentration to single-cell area. The line represents the mean YAP/TAZ concentration in each size range. The error margin corresponds to one standard deviation in the same size bin. Concentrations have been normalised to the means across all sizes for viewability. For each cell line, the relationship is shown across the size range: 0.5*mean – 2* the mean cell size.

We modelled the concentrations of YAP/TAZ (integrated intensity/cell area, mean intensity) as power law relationships with cell area, [*Y AP/T AZ*] = *aA*^*b*^, such that we could define a ‘scaling factor’, ‘b’, for each species (Fig. 1G). In a log-log plot, ‘b’ is given as the gradient and log(a) is the y-intercept. Negative values of ‘b’ correspond to the dilution of the protein with increasing cell size (sub-scaling), 0, linear scaling, and positive values, concentration of the protein with increasing size (super-scaling) (Fig. 1H). Fitting ‘a’ and ‘b’ values to each cell line’s F-actin concentration profile, we observed linear scaling between cell size and F-actin (b ranging from -0.2 to 0.2, (Fig. 1I, Supp.Table 2) as observed in previous studies (4). However, when applying the same analysis to whole cell YAP/TAZ concentration, we observed, for all cells lines, a clear sub-scaling relationship between cell size and whole cell YAP/TAZ concentration (b -0.35 - -0.65), indicating that whole cell YAP/TAZ dilutes as a cell gets larger (Fig. 1I, Supp.Table 3).

To investigate the decrease in whole cell YAP/TAZ concentration with cell size, we also analysed the abundance (integrated whole cell intensity, rather than mean) -size relationship and observed that the whole cell YAP/TAZ abundance increases with cell size, but not at a rate sufficient to maintain a constant concentration (b 0.6). Total YAP/TAZ increased with size at all sizes implying continued net-synthesis (That is, synthesis must be outpacing degradation, Supp.Table 4, Supp. Fig. 2). F-actin exhibited 1:1 abundance scaling indicating synthesis exactly offsets the effect of cell size on concentration. (Supp. Fig. 2, Supp.Table 5).

**Figure 2.**
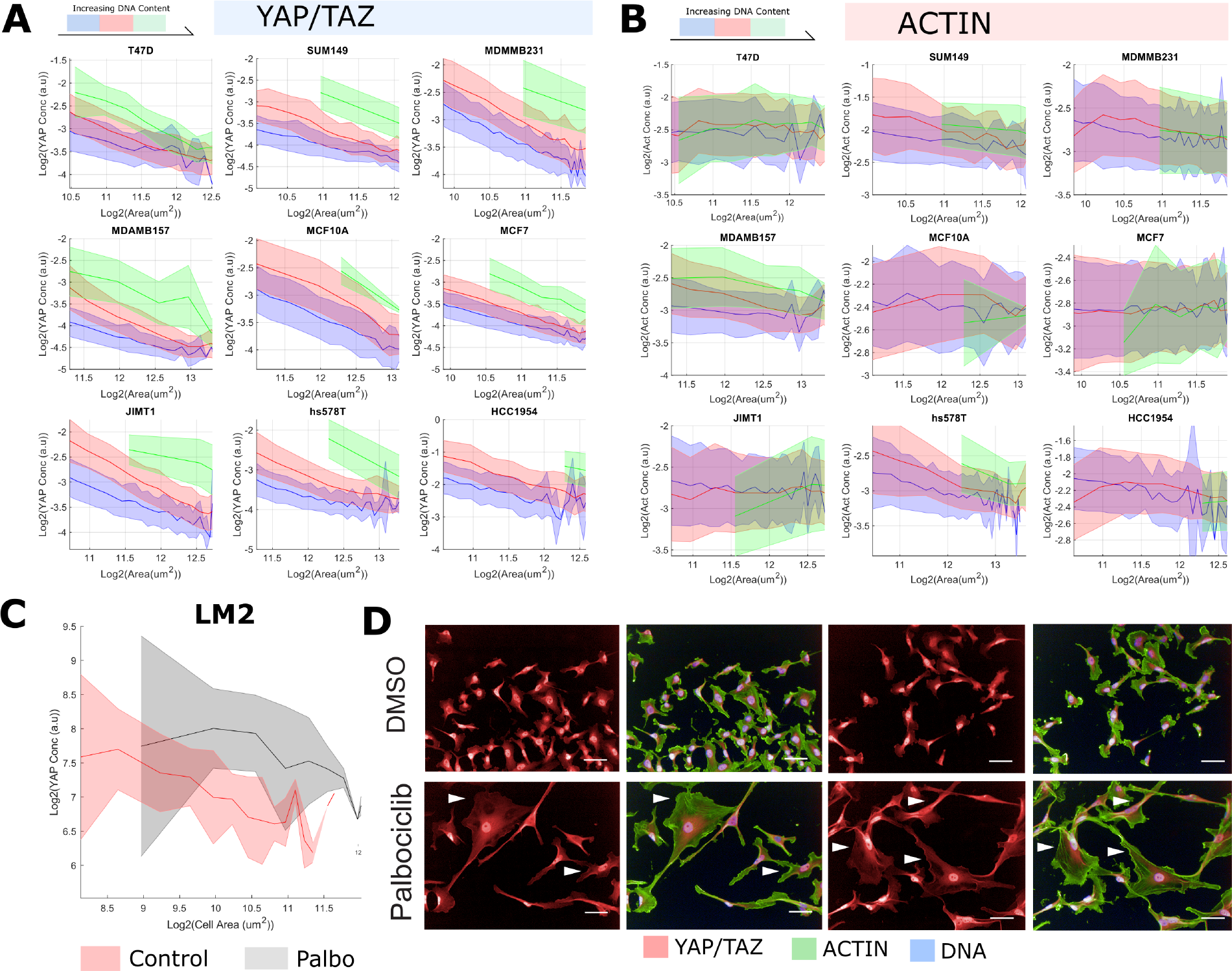
YAP/TAZ concentration, but not scaling, is sensitive to DNA-content and cell cycle progression A) Log-log plots relating YAP/TAZ concentration to single-cell area across DNA bins (blue the least DNA, red, green, yellow the most). Cell membership to each ‘bin’ was determined by kmeans clustering on the DNA content. The shaded area denotes one standard deviation of the cell size distribution about that size bin. B) Log-log plots relating Actin concentration to single-cell area across DNA bins (blue the least DNA, red, green, yellow the most). Cell membership to each ‘bin’ was determined by kmeans clustering on the DNA content. YAP/TAZ-size scaling is sensitive to DNA content while Actin scaling, is not. The shaded area denotes one standard deviation of the cell size distribution about that size bin. C) Dilution of YAP/TAZ with increasing cell size in LM2 breast cancer cells. The line represents the mean YAP/TAZ concentration in each size range. The error margin corresponds to one standard deviation in the same size bin. The relationship in untreated cells is shown in red, that for Palbociclib treated cells in grey. D) Representative images demonstrating the effects of palbociclib treatment on YAP/TAZ abundance and translocation in LM2 cells. Scale bar denotes 50um, YAP/TAZ in red, actin in green, DNA stain in blue.

To ensure that YAP alone was diluting with size, and that the measured effect was not an artefact generated by the erroneous recognition of TAZ by the antibody, we repeated the experiment with specific YAP and TAZ antibodies (36). In both cases, we observed dilution of whole cell YAP or TAZ protein with increasing cell size relative to F-actin (Supp.fig. 1). Together, these data reveal that both whole cell YAP and TAZ dilute with increasing cell size. As YAP/TAZ increases in abundance with as cells enlarge, this dilution is not due to net degradation of YAP/TAZ at larger cell sizes, but rather the effect of an expanding volume acting on an insufficient synthesis-degradation balance.

### B. YAP/TAZ concentration, but not scaling, is sensitive to DNA-content and cell cycle progression

As YAP/TAZ levels did not correlate with size across lines, but did so within lines, we hypothesised that sub-scaling of whole cell YAP/TAZ may be due to growth during cell cycle processes. To investigate this, within each line, we initially binned cells based on their DNA content (integrated Hoechst intensity). Bin sizes were constrained such that each bin centroid represents double the value of the preceding bins. Conducting the previous analysis on each DNA ‘bin’ within each line, we noticed that while the mean whole cell YAP/TAZ concentration at any given size increased for each doubling of the cell DNA (each DNA content ‘bin’, 1.3-1.6 factor increase) (Supp.Table 3, Fig. 2A), the scaling factor ‘b’ showed no obvious dependence on the amount of DNA. We performed the same analysis for the F-actin concentration and noticed no relationship between ‘b’ or mean concentration and DNA content (Fig. 2B, Supp.Table 2).Whole cell YAP/TAZ and F-actin abundance showed a consistent positive scaling factor (0.4-0.6) across all DNA contents (Supp.Table 4/5, Supp.fig 4/5).

As whole cell YAP/TAZ dilutes within the first and second DNA ‘bins’, loosely approximating ‘G1’ and ‘G2’, and undergoes a DNA-correlated regeneration between the two, potentially at S-phase, we sought to more rigorously investigate the relationship between cell cycle progression and YAP/TAZ concentration. We stained MCF10A cells for YAP/TAZ, PCNA and CCNA2 and trained a linear classifier to distinguish G0, G1, S and G2 cells using 110 CCNA2 and PCNA intensity features (methods) across 20,000 single cells. Conducting a scaling analysis within each stage, we observed that the negative size-YAP/TAZ concentration scaling is preserved across all stages besides G0 (Supp.fig. 6). Moreover, by binning the cells by area and calculating the mean YAP/TAZ concentration in each stage, we observed that the whole cell YAP/TAZ concentration increases from G1 to G2 and that smaller cells exhibit a greater whole cell YAP/TAZ concentration at each stage, further corroborating the previous analyses (Supp.fig. 6).

To test the idea that whole cell YAP/TAZ dilution relates to cell cycle progression, we assessed YAP/TAZ levels in Palbociclib treated LM2 cells. Palbociclib arrests cells at the G1/S transition by inhibiting CDK4/6 activity (40). We found that the average whole cell YAP/TAZ concentration was unchanged in Palbociclib treated cells despite the two-fold increase in size. YAP/TAZ dilution, however, still occurred with increasing size in the Palbociclib treated population (Fig. 2C/D). Together, these data show that the regulation of the cytoplasmic YAP/TAZ concentration is closely tied to the cell cycle, with dilution only being observed in cycling cells, and YAP/TAZ synthesis being strongly upregulated around S-phase.

### C. A constant nuclear concentration of YAP/TAZ is maintained across cell sizes despite whole cell dilution

Having observed a sub-scaling relationship between whole cell YAP/TAZ and cell size, we were interested in how this related to the nuclear translocation and concentration of YAP/TAZ. We quantified the N/C ratio of YAP/TAZ at the single cell level, for each cell line. Interestingly, YAP/TAZ ratio increased with increasing cell size, exhibiting the opposite relationship to (whole cell) YAP/TAZ concentration. The N/C ratio of YAP/TAZ also changed with DNA content, where the mean YAP/TAZ ratio, decreased across each DNA ‘bin’. We noted that ‘b’, the scaling factor between cell area and N/C, remained unchanged across DNA contents for all cell lines (Fig. 3A, Supp.Table 6).

**Figure 3.**
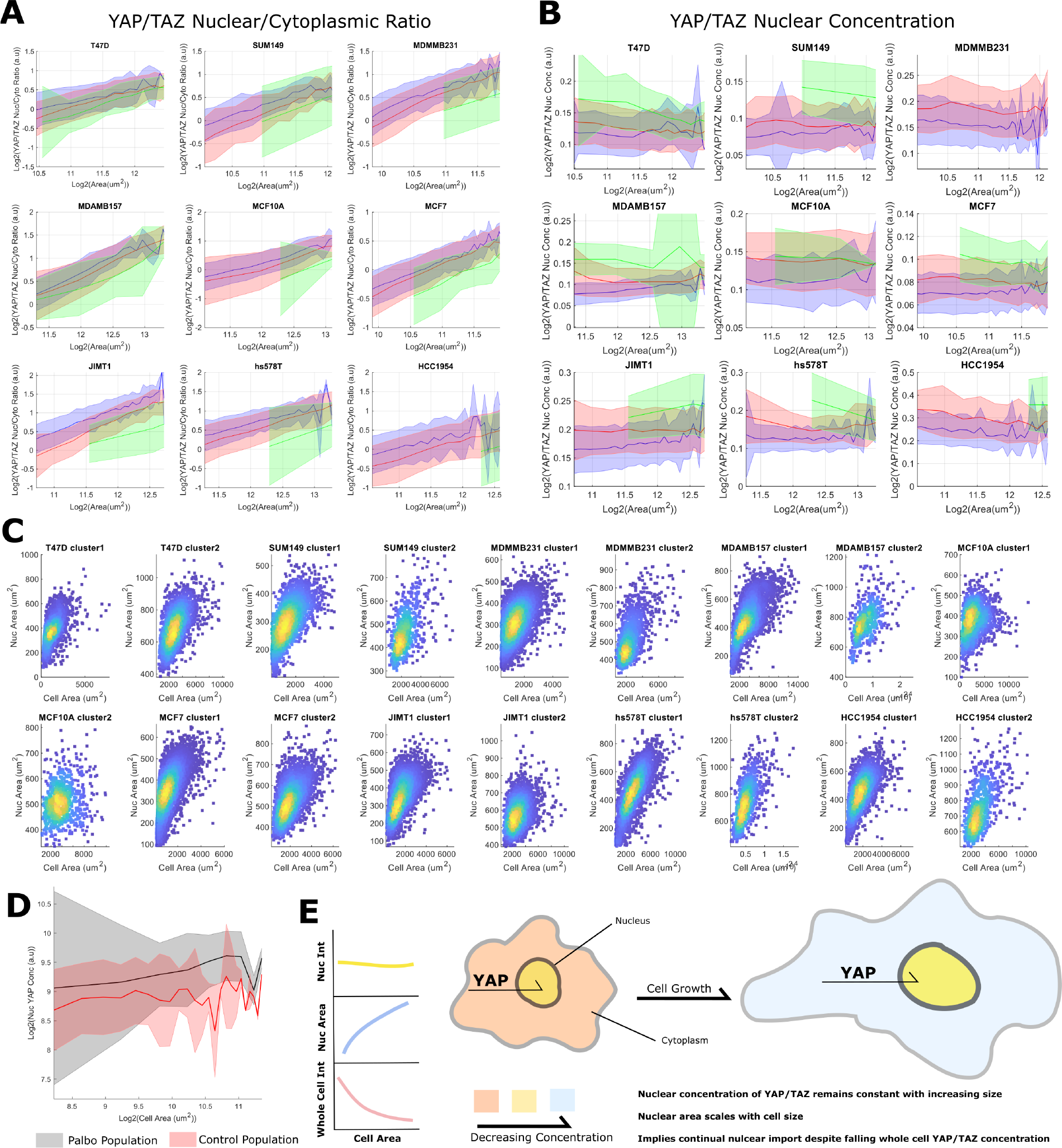
A constant nuclear concentration of YAP/TAZ is maintained across cell sizes despite whole cell dilution: A) Log-log plots relating YAP/TAZ nuclear-cytoplasm ratios and single cell area across lines and each DNA content bin (as determined by kmeans clustering on the integrated Hoechst intensity and nuclear area). Blue represents the lowest DNA content, then red, green and yellow, the most. In all cases, YAP/TAZ ratio positively scales with cell area but decreases with increasing DNA content. The shaded area denotes one standard deviation of the cell size distribution about that size bin. B) Nuclear YAP/TAZ, single cell area relationship plotted across DNA content bins for each cell line. The shaded area denotes one standard deviation of the cell size distribution about that size bin. C) Relationship between cell and nuclear areas across high and low DNA content clusters for each cell line (cluster 1 has less DNA than cluster 2). Nuclear and cell areas continually scale within DNA bins and is not related to DNA synthesis alone. D) The effect of Palbociclib on the nuclear YAP/TAZ concentration. Palbociclib does not affect scaling behaviour but increases the total nuclear concentration. E) A cartoon summarising the major findings of the section: Nuclear YAP/TAZ concentration is constant across sizes, nuclear area scales with size, together implying continual YAP/TAZ nuclear import despite a falling whole cell concentration.

**Figure 4.**
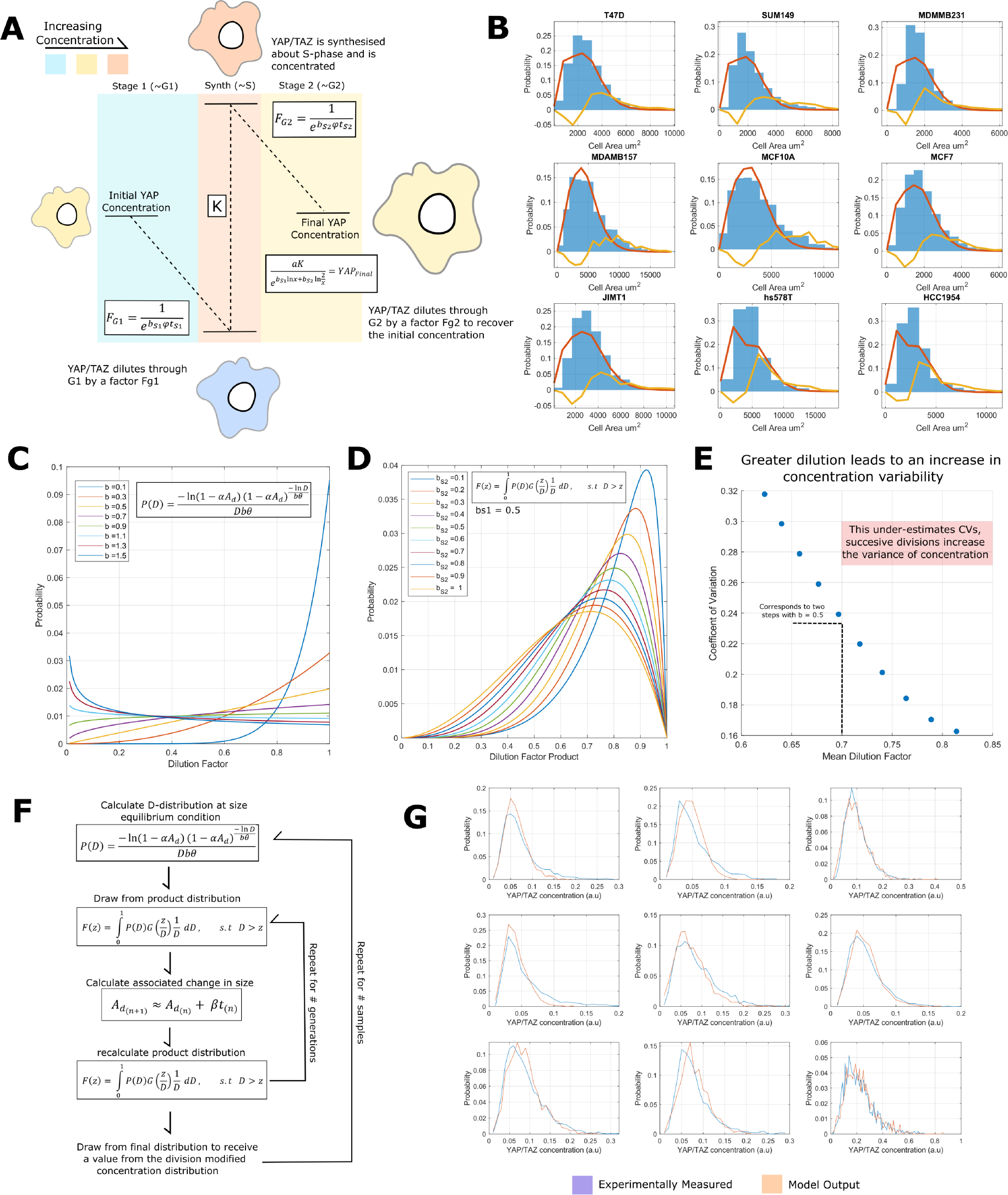
YAP/TAZ scaling behaviour is consistent with a dilution-synthesis-dilution scheme. A) Schematic describing the logic of the dilution-synthesis-dilution system. Relevant dilution factors are marked on each stage. The colours correspond to the final concentration at the end of the stage. Blue is low, orange high. B) Model fits on the single cell ‘G2’ (as determined by kmeans clustering on the integrated Hoechst intensity) size distributions, blue represents experimental data, orange is the calculated distribution and yellow is the Kullbeck-Leibler Divergence between the distributions. C) The dilution factor probability distribution (P(D)) across a range of scaling factors. D) The dilution factor product probability distribution F(z) across a range of scaling factors. E) The relationship between the coefficient of variation and the mean of F(z). The CV increases as the mean decreases. F) Schematic describing how the dilution factor distribution P(D) is used to calculate the expected YAP/TAZ concentration distribution G) Measured YAP/TAZ intensity distributions (blue) and the associated model fits (orange). In all cases we observe excellent agreement.

We next investigated the scaling of the nuclear concentration of YAP/TAZ cell area. Strikingly, the nuclear concentration distribution of YAP/TAZ was almost entirely insensitive to increases in cell size, exhibiting a constant mean and variance across all measured areas. Increases in the DNA content did increase the average nuclear YAP/TAZ level, but not sufficiently to maintain the same YAP/TAZ ratio across DNA bins (Fig. 3B, Supp.Table 7). Importantly, as nuclear and cell size correlate (Fig. 3C), even without an increase in DNA content, this result implies continual transport of YAP/TAZ into the nucleus as cells grow such to maintain a constant concentration distribution across differently sized cell populations. Thus, nuclear transport of YAP/TAZ is coupled to cell size in order to maintain a steady-state level of YAP/TAZ as the cytoplasmic pool becomes diluted.

To determine if nuclear concentration was also dependent on CDK4/6 activity, we analysed nuclear YAP/TAZ levels in Palbociclib treated cells. We observed that while the nuclear concentration was invariant to cell size, it was sensitive to Palbociclib treatment; as treated cells exhibited higher nuclear YAP/TAZ concentrations than control cells. (Fig. 3D). Thus CDK4/6 activity and/or cell cycle progression is necessary to couple nuclear transport of YAP/TAZ to cell size.

Together, these data suggest that while the concentration of the cytoplasmic pool of YAP/TAZ is a function of cell size and volume, the nuclear YAP/TAZ concentration is regulated independently. As a transcriptional regulator, this implies that as cell divide they maintain a constant pool of nuclear YAP/TAZ activity despite falling cytoplasmic concentrations. Indeed, this is particularly striking given the strong correlation between the nuclear and whole cell YAP/TAZ concentration (Fig. 1.D/E). Given the scaling between nuclear and cell area, necessitating continual import of YAP/TAZ, this may be driven by nuclear transport machinery (summarised in Fig. 3E).

### D. Integration of YAP/TAZ size-scaling and stochastic cell division determines YAP/TAZ heterogeneity

The previous analyses are consistent with a scheme by which a constant amount of YAP/TAZ is maintained through dilution in G1, synthesis at S/G2 and a further dilution through the subsequent G2, before the inheritence of YAP/TAZ by daughter cells (A dilution-synthesis-dilution, or DSD, scheme (Fig. 4A)). We sought to understand how such a system would behave across division cycles and what consequences this could have for the cell population. To understand how YAP/TAZ scaling affects the YAP/TAZ concentration distribution, we integrated a DSD model with a simple cell cycle system (a two-stage system exhibiting adder size behavior) (9). This model assumes linear dependencies between cell area and the probability to advance in the cell cycle stage and cell area and growth rate. The cell area probability distribution under these constraints is given as a hypo-exponential function *J* (*A, S, n*):

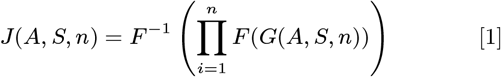

where:

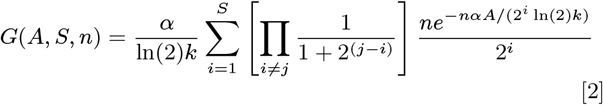

Where: ‘Ad’ is the cell area at division, ‘a’ is a proportionality constant between Ad and cell cycle advancement probability, ‘k’ a proportionality constant between Ad and growth rate, ‘S’ is the number of divisions that have occurred, and ‘n’ is the number of cell cycle ‘stages’ per division. ‘F(f(x))’ denotes the Fourier transform of f(x), and *F* ^*−*1^, the inverse. (See methods (calculating area distributions) and supplemental information for details).

To ensure the applicability of this model, we fit equ. 1 to the (G2) cell area distributions of our cell lines using a two-stage system and found good agreement (methods). Here ‘stage’ refers to the number of growth stages in a cycle (Fig. 4B). Indeed, the CVs of the distributions lie well within the predicted bounds of a two-stage system 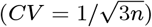. For *n* = 2, 0.4 *< CV <* 0.57 (Supp. Table 7) (9).

Expressing the ‘cell cycle advancement time’ (time spent in each cycle stage, CCA) distribution in terms of the dilution factor ‘D’ (*D* = 1*/* exp(*bkt*), where ‘b’ is the scaling factor, ‘k’ the proportionality constant between size and growth rate, and ‘t’ is time) gives (Fig. 4C) (see supplemental information):

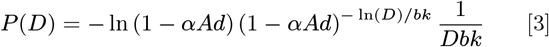

As two dilution events happen in sequence, we were interested in the product distribution of *P* (*D*) and a second dilution event, denoted as *F* (*z*) (Fig. 4D):

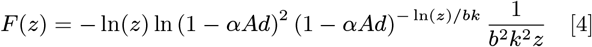

Where *z* is the product of two random ‘D’ variables distributed according to equ. (3). For simplicity, we considered the two sequential dilution events to be identical, such that the cell grows equally across G1 and G2 and exhibit the same scaling behavior. From *F* (*z*), we could calculate the coefficient of variation (*σ/σ*):

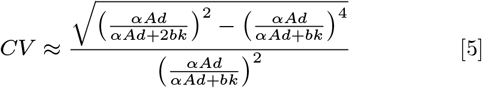

Which approximately linearly scales with ‘b’ for 0 *< b <* 1 (Fig. 4E); note we have expanded the mean and variance about *a* = 0 to obtain equ. (5) (*α* values do not exceed 10^*−*5^) (methods). Correlating scaling factors against the CV of YAP/TAZ intensity distributions, we observed a strong negative trend for 7*/*9 of our cell lines (Supp.fig. 8).

It is important to note that average ‘b’ values are not the only source of YAP/TAZ variance in this system; differences in size regulation and the area dependence of ‘K’ and ‘b’ all contribute to YAP/TAZ variability (Supp.fig. 7). Indeed, the MDA-MB-231 and JIMT1 cell lines, those with high scaling factors (*b*≈0.54) but comparatively low YAP/TAZ variance (*CV*≈0.37), have more homogeneous size distributions than most of the other cell lines (Supp. Table 8, *CV* ≈ 0.48) and are some of the few to exhibit an approximately constant ‘K’ value (Supp.fig. 7) providing an explanation for their departure from a linear scaling factor-variance relationship.

**Figure 5.**
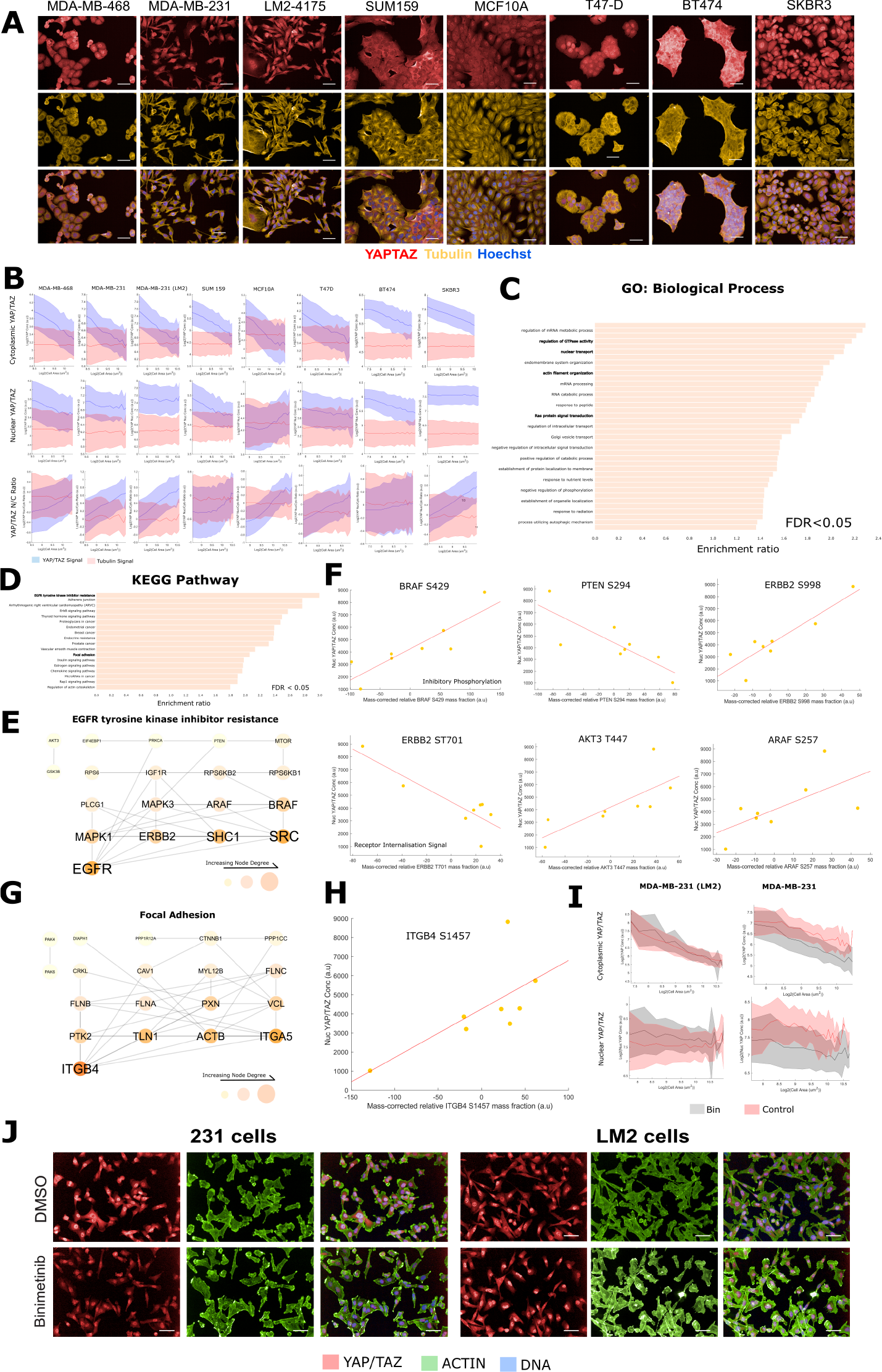
The nuclear YAP/TAZ concentration distribution is associated with altered RAS, adhesion and nuclear transport signalling processes: A) Representative images of the 8-lines across which we conducted phospho-proteomic experiments. Scale bar denotes 50um, YAP/TAZ in red, tubulin in yellow, DNA stain in blue. B) Recapitulation of whole cell YAP/TAZ dilution with increasing cell size and perfectly scaling nuclear concentration against a tubulin standard in a separate experiment and cell line panel. The central line denotes the mean YAP/TAZ cytoplasmic concentration (TOP), nuclear concentration (MID) or N/C ratio (BOT) in each size bin. The error bars correspond to one standard deviation in that bin. Tubulin signal is shown in red and YAP/TAZ in blue. C) Themes from the biological process noRedundent dataset enriched in the list of phosphopeptides most predictive of a cells nuclear YAP/TAZ concentration. All enrichments are significant to *FDR <* 0.05. D) Themes from the KEGG pathway dataset enriched in the list of phosphopeptides most predictive of a cells nuclear YAP/TAZ concentration. All enrichments are significant to *FDR <* 0.05. E) A network of the interacting members of the phosphopeptides predictive of the nuclear YAP/TAZ concentration under the ‘EGFR tyrosine kinase inhibitor resistance’ KEGG pathway. Interactions were derived from the STRING database, only experimentally determined physical interactions are shown. Node size, label size and colour are proportional to the node degree. F) Example relationships between the nuclear YAP/TAZ concentration and enriched phosphopeptides from the ‘EGFR tyrosine kinase inhibitor resistance’ KEGG pathway. G) A network of the interacting members of the phosphopeptides predictive of the nuclear YAP/TAZ concentration under the ‘Focal Adhesion’ KEGG pathway. Interactions were derived from the STRING database, only experimentally determined physical interactions are shown. Node size, label size and colour are proportional to the node degree. H) Example relationships between the nuclear YAP/TAZ concentration and enriched phosphopeptides from the ‘Focal Adhesion’ KEGG pathway. I) The effect of Binimetinib treatment on YAP/TAZ whole cell and nuclear scaling in LM2 and 231 cells. Binimetinib increased the nuclear concentration of YAP/TAZ across all sizes in LM2 cells. Binimetinib had the opposite effect in M231 cells. J) Representative images demonstarting the effects of binimetinib treatment on YAP/TAZ abundance and translocation. Scale bar denotes 50um, YAP/TAZ in red, actin in green, DNA stain in blue.

**Figure 6.**
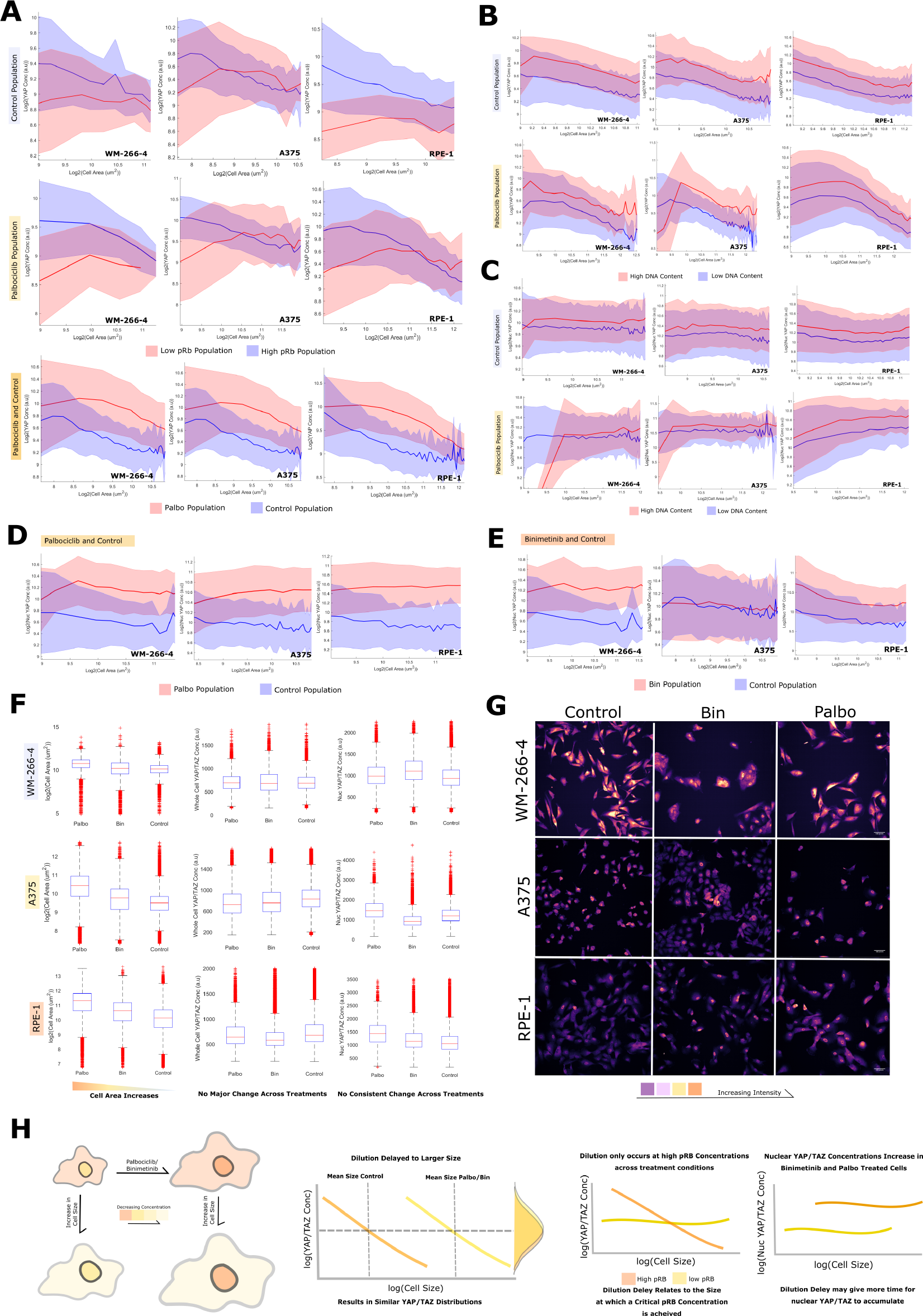
YAP/TAZ dilution behaviour is conserved across melanoma and RPE cells: A) YAP/TAZ concentration – size relationship across WM266-4, A375 and RPE1 cells. Top shows whole-cell YAP/TAZ size scaling in the high (blue) and low (red) phospho-RB1 populations in untreated cells. The middle shows the same in the Palbociclib treated context. The bottom directly compares YAP/TAZ size scaling in the high pRB1 populations in the Palbociclib treated and control cells. In all cases, the middle line represents the mean value of cells about that size ‘bin’. The width of the shaded area relates to the standard deviation of cells in that same bin. B) The relationship between YAP/TAZ dilution and DNA content across control (top) and Palbociclib treated (bottom) cells. The high DNA content cluster is in red, the low cluster in blue. C) Nuclear YAP/TAZ concentration against cell size across DNA content in control (top) and Palbociclib treated (bottom) cells. The high DNA content cluster is in red, the low cluster in blue. D) Nuclear YAP/TAZ across cell sizes compared across low DNA cluster Palbociclib treated (red) and control (blue) cells. In all three cell lines, Palbociclib increased the nuclear concentration of YAP/TAZ across all cell sizes. E) Nuclear YAP/TAZ across cell sizes compared across low DNA cluster Binimetinib treated (red) and control (blue) cells. In WM266-4 and RPE1 cells, Binimetinib increased the nuclear concentration of YAP/TAZ across all cell sizes. It had no effect in A375 cells. F) Boxplots summarising the effects of Palbociclib and Binimetinib at the population level on; cell area (left), whole cell YAP/TAZ (middle) and nuclear YAP/TAZ (right). G) Representative images of YAP/TAZ signal across cell lines in each treatment. Colour is proportional to the YAP/TAZ intensity. H) Cartoon summarising the major findings of this section: Palbociclib/Binimetinib do not reduce whole-cell YAP/TAZ despite increasing cell size because scaling is delayed to a larger size, this appears related to the phosphorylation of RB1 in these cell lines, the delay may give more time for nuclear YAP/TAZ to accumulate.

**Figure 7.**
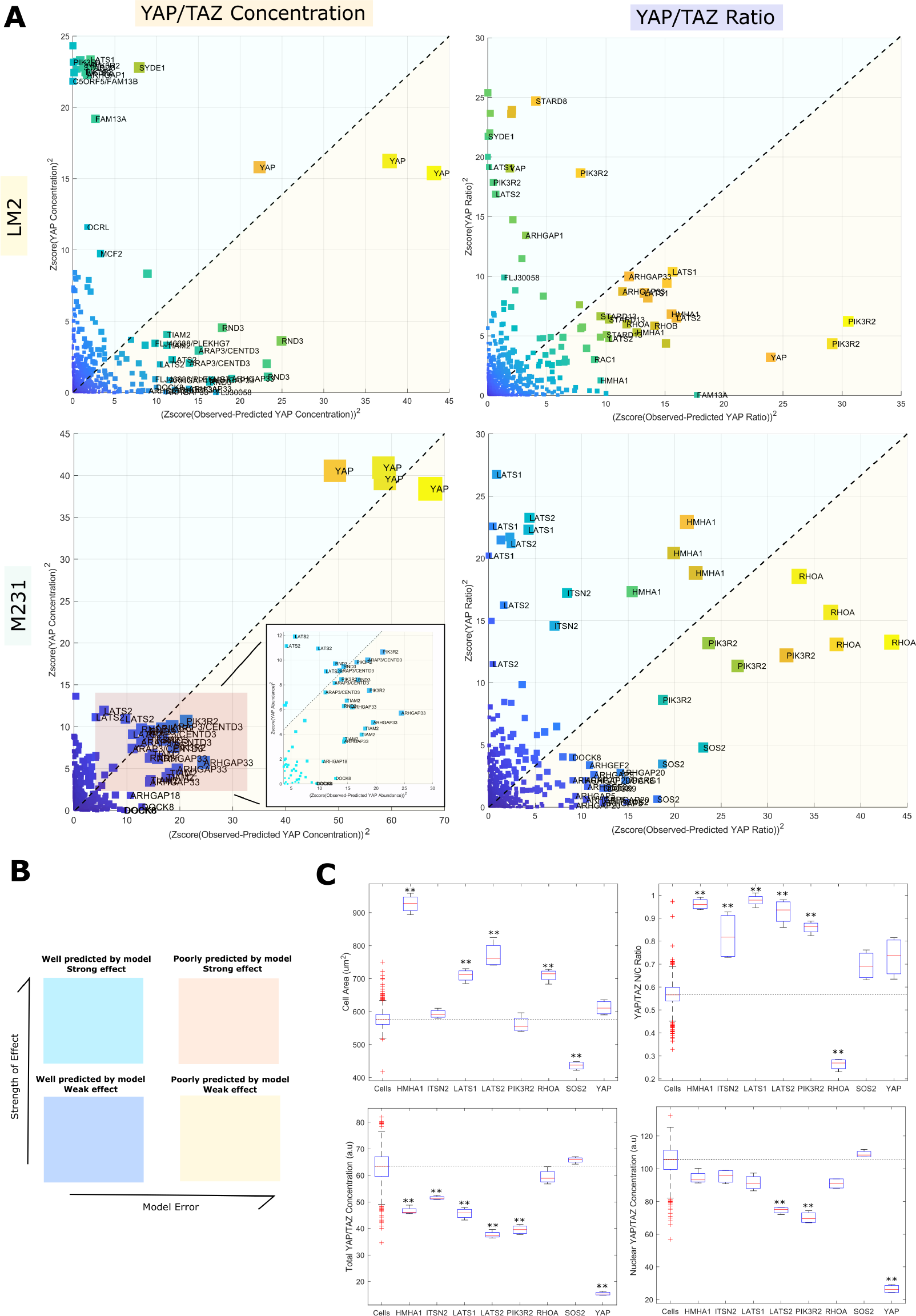
Dilution of cytoplasmic YAP/TAZ drives increasing nuclear/cytoplasmic ratios following gene depletion A(top left)) The relationship between the squared YAP/TAZ concentration (Z-score) and squared model error (Z-score) for each knockdown state in LM2 cells. The colour and size of each datapoint corresponds to the distance from the origin. A(bottom left)) The relationship between the squared YAP/TAZ concentration (Z-score) and squared model error (Z-score) for each knockdown state in M231 cells. The colour and size of each datapoint corresponds to the distance from the origin. Inset: zoom in on the red marked region. We note a strong overlap in hits across the two cell lines. A(top right)) The relationship between the squared YAP/TAZ ratio and squared model error for each knockdown state in LM2 cells. KDs affecting YAP/TAZ ratio are not redundant with those affecting concentration. The colour and size of each datapoint corresponds to the distance from the origin. Inset: zoom in on the red marked region. A(bottom right)) The relationship between the squared YAP/TAZ ratio and squared model error for each knockdown state in M231 cells. The colour and size of each datapoint corresponds to the distance from the origin. Inset: zoom in on the red marked region. B) A guide to the regions of the graphs shown in ‘A’. C) Comparisons of YAP/TAZ concentration, YAP/TAZ nuclear concentration, YAP/TAZ N/C ratio and cell area across knockdown states by an N-way ANOVA test (n = 200 for control, 4 for all else, ** denotes *P <* 0.01).

To capture the concentration distribution of whole cell YAP/TAZ, we simulated equ. 4 across multiple generations (Fig. 4F) (methods). Parameter (*Q*) values (*bs*1, *bs*2, *K*) were taken from the size-Q curves for each cell line. Comparing the predicted distributions to those measured, we observed excellent agreement showing that, in these cell lines, the concentration distribution of whole cell YAP/TAZ can be explained by size-dependent stochastic cell division acting on a dilution-synthesis-dilution system (Fig. 4G). Thus, coupling of YAP/TAZ nuclear transport to size is important to suppress variability in YAP/TAZ levels over successive generations.

### E. The mean nuclear YAP/TAZ concentration across sizes is associated with altered RAS, adhesion and nuclear transport signalling processes

Having observed the dilution of cytoplasmic (and whole cell) YAP/TAZ with increasing size, and that size had no tangible effect on the mean nuclear concentration (or concentration distribution across the population), we were interested in how continual import could be sustained across sizes whilst the cytoplasmic pool depletes. To investigate this, we combined high-throughput imaging and phosphoproteomic experiments across a separate panel of eight cell lines (semi-redundant with the previous panel) (Fig. 5A, Table.1). The cell lines selected were similar sizes (within a 2-fold range) to prevent size-related phosphorylation events colouring the investigation of nuclear YAP/TAZ and ratio correlates. YAP/TAZ exhibited size sub-scaling behaviour at the whole cell level, but not in the nucleus (relative to tubulin intensity), as is consistent with the prior dataset (Fig. 5B).

We predicted the cell lines nuclear YAP/TAZ and ratio from the phosphoproteomic expression data using partial-least squares regression (PLSR). For this, the expressions of each phosphopeptide were ‘corrected’ such that they reflected how much more/less expressed they were than expected given the detected expression of the unphosphorylated peptide (see methods). This eliminated the trivial correlation between proteomic and phosphoproteomic data and gives information on the signalling state of the cells. From the PLSR model, we could calculate the contribution of each phospho-peptide and thus, how predictive each phosphopeptide was, as achieved through calculation of a ‘Variable importance to projection’ (VIP) score.

Mean nuclear YAP/TAZ concentrations were predicted by mass corrected expressions of phosphopeptides enriching for: RTK/MAPK signalling (KEGG pathway ‘EGFR tyrosine kinase inhibitor resistance’, FDR *<* 0.05). These included several core regulators of the MAPK pathway including:

ERBB2 (HER2) (T701), ARAF (S269), SRC (S75), SOS1 (S1178), MAPK1 (T185), MAPK3 (T202), PTEN (S294) and MTOR (S1261) (Fig. 4C/D). We also observed a clear association between nuclear YAP/TAZ concentration and focal adhesion signalling (KEGG pathway ‘Focal adhesion’, *FDR <* 0.05), with phosphorylations on TLN (Y70), VCL (S346), PXN (S533), PTK2 (S29), PAK4 (S474), PAK6 | | (S616), ITGB4 (S1457/4) strongly correlating (│*R*│ > 0.7) (Fig. 5C/D/E/F/G/H). Together, these sites suggest that enrichment of nuclear YAP/TAZ is related to ERK/MAPK signalling activation and the maturation of focal adhesions (41).

Strikingly, when predicting the YAP/TAZ N/C ratio, we observed differential phosphorylation on multiple regulators of nuclear transport across cell lines with high and low YAP/TAZ ratios (GO:0051169, ‘nuclear transport. *FDR <* 0.05) These included the nucleoporins, NUP133/153/210/35/188/85 and NDC1 (S406), LMNA (S403), the RAN binding proteins RANBP2 (S2280) and 3 (S27), and XPO1 (S1055), a protein recently directly implicated in YAP/TAZ export from the nucleus (42).

To investigate the role of EGFR/MAPK signalling in YAP/TAZ translocation, we treated two breast lines, MDA-MB-231 and MDA-MB-231-LM2, with Binimetinib, a MEK inhibitor. In LM2 cells, Binimetinib treatment resulted in an increase in nuclear YAP/TAZ per cell size whilst having no obvious effect on the scaling of the whole cell YAP/TAZ concentration implying increased translocation. Conversely, in 231 cells, Binimetinib partially reduced nuclear and whole cell YAP/TAZ levels (Fig. 5I/J), however, increased the N:C ratio, as in LM2 cells implying increased nuclear import. Taken together these data suggest that RTK-MAPK signalling couples cell size to YAP/TAZ nuclear translocation. Inhibition of MEK signalling disrupts the coupling leading to changes in nuclear translocation.

### F. YAP/TAZ dilution behaviour is conserved

After thoroughly characterizing the dilution behaviour of YAP/TAZ in normal and cancerous breast cells, our study was expanded to examine this effect in different cell contexts and assess its generality as a phenomenon. Specifically, we conducted imaging experiments on WM-266-4 and A375 melanoma cells, as well as retinal pigment epithelial cells (RPE-1). The objectives were to investigate whether: 1) Whole cell YAP/TAZ dilutes as cells enlarge, 2) the nuclear YAP/TAZ concentration remains constant with increasing size, and 3) CDK4/6 and MEK inhibition promote nuclear accumulation of YAP/TAZ.

Strikingly, across all three of the added lines, YAP/TAZ dilution was conserved. To more formally test the relationship between YAP/TAZ dilution and the cell cycle, we also stained these lines for pRB1. Interestingly, dilution only occurred in cell populations with high pRB1 in these lines, corroborating the cell cycle dependency seen in the breast cells (Fig. 6A), and the absence of YAP/TAZ dilution in G0 cells. This extended to the Palbociclib treated population, although, cells with higher pRB1 tended to larger in this setting, delaying dilution to a larger cell size (Fig. 6A). The YAP/TAZ concentration, but not the scaling behaviour, was found to be sensitive to DNA content in all cell lines in both the control and Palbociclib treated populations (Fig. 6B).

As in breast cells, the nuclear concentration of YAP/TAZ remained constant with cell size in the background of whole cell dilution, and this was similarly found sensitive to the DNA content of the cell (Fig. 6C). Palbociclib increased the nuclear concentration of YAP/TAZ per cell size in all cases (Fig. 6D). Binimetinib exhibited similar behaviour in Wm-266-4 and RPE-1 cells, but failed to elicit a response from A375 cells (Fig. 6E).

At a population level, both Palbociclib and Binimetinib increased average cell size, with Palbociclib having a stronger effect (Fig. 6F). Despite this, neither treatment had any obvious effect on the whole cell YAP/TAZ concentration. This is presumably due to the delay of dilution in either treatment (Fig. 6F). Palbociclib increased the average nuclear YAP/TAZ concentration in all cases. Binimetinib treatment echoed the result in WM-266-4 and RPE-1 but interestingly reduced nuclear YAP/TAZ in A375 cells (Fig. 6F). The results of this section are summarised in Fig. 6G. Together, these data show that the YAP/TAZ dilution phenomenon extends to the melanoma and RPE cell contexts evidencing the generality of the effect.

### G. Dilution of cytoplasmic YAP/TAZ drives increasing nuclear/cytoplasmic ratios following gene depletion

Our integrated analysis suggested that RTK-RAS-ERK, focal adhesions, and nuclear transport are key processes which couple cell size to YAP/TAZ nuclear transport and act as mechanism to maintain steady state levels of nuclear YAP/TAZ as cells grow. To identify additional factors that may act to couple YAP/TAZ nuclear transport to cell size we performed genetic screen where we systematically depleted 82 RhoGEFs, 67 GAPs, and 19

Rho GTPases, across 300,000 LM2 and MDA-MB-231 cells (39). We focused on RhoGEF, RhoGAPs, and RhoGTPases as these are well-established regulators of both YAP/TAZ and cell morphology - thus are excellent candidates for genes that may act to couple nuclear transport of YAP/TAZ to cell growth. We also included in this screen siRNAs that targeted YAP and TAZ, as well as components of the Hippo pathway such as LATS1 and LATS2.

Identifying regulators of YAP/TAZ transport following perturbation is complicated by the fact that any given gene could potentially regulate YAP/TAZ concentration by affecting volume, by affecting signalling that regulates transport, or both. Conducting a scaling analysis on the untreated cells (LM2 and MDA-MB-231 cells) (*n*≈ 80, 000), we observed the same whole cell YAP/TAZ concentration sub-scaling behavior as in the prior analyses (*b*≈0.35). Similarly, the nuclear concentration distribution remained approximately constant with size, leading to an increase in average YAP/TAZ n/c ratio driven by population-level cytoplasmic dilution.

Gene depletions did affect the nuclear and cytoplasmic concentration distribution of YAP/TAZ. For example, depletion of genes such as SPATA13, RALBP1, and BCR increased average YAP/TAZ concentrations/n/c ratios for any given cell size in MDA-MB-231 cells. Investigating more deeply, we constructed partial least squares regression (PLSR) models which predicted YAP/TAZ concentration and ratio in control cells as functions of 114 measured morphological and intensity features (methods) (Supp.fig. 11/12). Calculating variable importance to projection (VIP) scores from the model, we observed that cell area (and associated correlates e.g., Nuclear area, eccentricity, etc.) most contributed to the prediction of YAP/TAZ concentration and n/c ratio (*V IP scores >* 1 are considered major contributors to the model). We applied this model to all treatment conditions (siRNA knockdowns) finding that, even under perturbation, the size-YAP/TAZ n/c ratio and concentration relationship persisted (Supp.fig. 11/12). Thus, the scaling of YAP/TAZ levels to size appears rarely affected by perturbations which affect size, morphology, or Rho GTPase pathway activation. While many siRNAs are affecting the YAP/TAZ n/c ratio, often they are doing so through manipulation of cell morphology and dilution of cytoplasmic (and total) YAP/TAZ, rather than increasing the nuclear concentration. These included genes canonically associated with increased cell size, including ECT2 and RACGAP1, known to induce cytokinetic failures and polyploidization when depleted (53).

Using this framework, we could also investigate genes that disrupt the coupling between YAP/TAZ and size; those which are most poorly predicted by the model are those which most perturb the relationship between YAP/TAZ and cell morphology. The clearest ‘hits’ across both cell lines included YAP itself, the YAP regulatory kinases LATS1 and LATS2, as well as RHOA, HMHA1, and PIK3R2. Unique to LM2 cells were ARHGAP33 and STARD13 whilst SOS2 and ITSN2 perturbed the relationship only in MDA-MB-231 cells (Fig. 7A/B).

Focusing on MDA-MB-231 cells, LATS1/2 and HMHA1 interestingly exhibited a very similar behavior to that captured by the model, in that they exhibited an increased cell area and YAP/TAZ ratio, and decreased whole cell YAP/TAZ concentration (Fig) However, these KDs led to an increase in YAP/TAZ ratio beyond what would be expected from an increase in size alone. Furthermore, LATS1/2 KD led to a small decrease in the nuclear YAP/TAZ concentration (Fig. 7C). Together, we conclude these genes affect the cytoplasmic levels of YAP/TAZ both via control of morphology, size, and signaling mechanisms (Fig. 7C).

This phenomenon was not universal, however; several KD states altered the YAP/TAZ ratio independently of cell area, such as PIK3R2 and ITSN2 (Fig. 7C). Interestingly, the reduction in YAP/TAZ ratio observed in these cases was nevertheless associated with a loss of cytoplasmic rather than an increase in the nuclear YAP/TAZ concentration. Importantly, this shows that an increase in cell area is not the only way to achieve a reduction in cytoplasmic YAP/TAZ concentration in this system (e.g., Increased degradation). These genes could be involved in directly regulating YAP/TAZ biosynthesis and/or stability. Of our ‘hit’ genes that increased YAP/TAZ n/c and cell area, only RHOA depletion led to a decrease in YAP/TAZ ratio, driven solely by a canonical reduction in nuclear, rather than total, YAP/TAZ concentration (Fig. 7C). It is unclear whether RHOA depletion leads to an increase in YAP/TAZ synthesis per cell size, such to offset the effect of size scaling on the mean YAP/TAZ concentration, or whether it decouples cell area from cell volume, leading to an anomalously high spread area skewing the result. Indeed, such an effect may underpin the behavior of PIK3R2/ITSN2 KD.

Together these data show that YAP/TAZ size-scaling and concentration are remarkably robust to perturbations in RhoGTPase signalling, in that only the depletion of very few RhoGEF/GAPs disturbed YAP/TAZ in a way inconsistent with the concomitant change in cell morphology. However loss of the core Hippo effectors, LATS1/2, and a master contractility regulator, RHOA (amongst others) successfully altered the relationship between cell morphology and YAP/TAZ regulation. Stable expression of these genes may be vital to maintaining a constant nuclear YAP/TAZ concentration distribution, and therefore signal sensitivity, as a cell grows.

## 2. Discussion

Here we have shown that cytoplasmic YAP/TAZ are sub-scaling molecules across cell types; specifically, diluting in G1, undergoing a surge in synthesis near S-phase before diluting again in G2. This is not unique to YAP/TAZ; seminal work on size-scaling phenomena showed that RB1 (and the associated Whi5 in yeast) exhibits extremely similar behaviour. However, unlike YAP/TAZ, RB1 is not continually synthesised throughout the cell cycle/across sizes (’b’ abundance 0.15 vs 0.4-0.6 for YAP/TAZ) (4, 5, 15). The concentration of RB1 is, therefore, more directly controlled by changes in cell volume, befitting of its putative role as a size-sensor, whereas the YAP/TAZ concentration is complicated by biosynthetic regulation.

That nuclear YAP/TAZ concentration distribution did not change across cell size bins suggesting that YAP/TAZ signalling is largely constant across small and large cells during proliferataion. That is, YAP/TAZ signalling is robust against changes in cell size occurring throughout a division cycle. Such robustness is not a rare phenomenon in biology, indeed, recent works developing models of biological signalling networks have observed remarkably low parameter sensitivity (54–57). A particularly striking example can be found in a model of the Drosophila segmentation network where, across 48 parameters and two orders of magnitude, if a parameter was assigned a random value, there was a 90% chance that it was associated with a functional network (58).

Amongst other mechanisms, a biological system may achieve robustness through adaptation (57). When investigating the signalling differences in cell lines with high/low average nuclear YAP/TAZ, we found that the expression of phosphopeptides relating to nuclear transport, adhesion and RTK-MAPK signalling best explained the differences, suggesting that these signalling systems may ‘adapt’ (are up/down regulated with increasing size over generations) to the depleting YAP/TAZ pool. Indeed, an increased activity of nuclear transporters (and decreased activity of exporters) with increasing size provides an intuitive explanation for how a constant nuclear concentration distribution, sustained by continual import, could be maintained under a falling cytoplasmic concentration. Indeed, XPO1 Ser1055, a phosphopeptide, upregulated in lines with a lower mean nuclear concentration, is an activated species known to control the export of YAP/TAZ (41, 42). Furthermore, conformation changes in nuclear pores have been shown to stimulate YAP/TAZ entry into the nucleus (59, 60). This can also be driven by increasing nuclear size and thus cell spreading and growth, imparting stress on the nucleus through cytoskeletal connections to the cell body, and may even be sufficient to sustain the nuclear YAP/TAZ concentration as the cell expands (61).

When perturbing MAPK and CDK4/6 activity, we observed an increase in the mean nuclear YAP/TAZ concentration per cell size. This may relate to YAP/TAZ’s role in prompting resistance to BRAF-MEK blockade. (62–67). As Binimetinib and Palbociclib exhibited similar effects, is it likely that MAPK’s role in promoting proliferation regulates nuclear YAP/TAZ. As neither treatment tangibly effected the scaling factor of the nuclear YAP/TAZ concentration, it is unlikely that MAPK signalling is dynamically regulated with increasing cell size to maintain robust signalling, but rather, determines the mean nuclear concentration to be maintained (and thus the nuclear concentration distribution across the population). Focal adhesion/mechanosignalling events may play a similar role. Indeed we did not observe any clear change in the ‘scaling factor’ of nuclear YAP/TAZ following depletion of many RhoGEFs and RhoGAPs which couple adhesion to signalling and morphogenesis, but only changes to the absolute quantity of YAP/TAZ per cell size.

Our theoretical model, integrating stochastic cell division with YAP/TAZ size scaling, revealed that more severe protein-size scaling results in a greater variance in the proteins concentration distribution; this may have drastic consequences for the cytoplasmic functions of YAP/TAZ; for example, YAP/TAZ has been shown to influence the spindle assembly checkpoint, potentially through its interactions with BUBR1 (68). Moreover, cytoplasmic YAP/TAZ is known to be a core component of the CTNNB1 destruction complex (69, 70). As YAP/TAZ and CTNNB1 co-operate as transcription factors in the nucleus (64, 71), this suggests that the cytoplasmic dilution of YAP/TAZ may also indirectly influence its nuclear activity in accordance with the putative importance of the YAP/TAZ nuc/cyto ratio (72–74).

Together, these data show that that YAP/TAZ can dilute as the cell increases in size. Remarkably, the nuclear concentration distribution is insensitive to the effect, demonstrating that cells have developed systems to mitigate the influence of protein dilution beyond just regulating their size. Such mechanisms may be crucial in overcoming the emergent heterogeneity associated with sub/super scaling behaviour across division cycles and for maintaining robust signalling throughout the cell cycle.

## Materials and Methods

### A. Cell Culture

The following human breast cell lines were investigated (novel in this study). T-47D and BT-474 were obtained from Nicholas Turner (ICR, London), SKBR3 cells were a kind gift from the laboratory of Olivia Rossanese (ICR), MDA-MB-468 cells were a kind gift from George Poulogiannis (ICR), MDA-MB-231 were obtained from Janine Erler (University of Copenhaguen, Denmark), LM2 cells (a highly metastatic subpopulation 4175 from MDA-MB-231, (38)) were obtained from Joan Massagu’
se (Sloan Kettering Institute, New York), while SUM159 were a kind gift from the laboratory of Rachel Natrajan (ICR). All the above cancer cell lines were grown in Roswell Park Memorial Institute (RPMI)-1640 culture medium (Gibco) supplemented with 10% heat-inactivated fetal bovine serum (FBS) and 1% penicillin/streptomycin. MCF10A cells were obtained from ATCC and were engineered to express endogenous mRuby-tagged PCNA (75). They were grown in DMEM/F12 supplemented with 5% horse serum, 10 μg/ml insulin, 20 ng/ml epidermal growth factor, 100 ng/ml cholera toxin, 500 ng/ml hydrocortisone, and 1% penicillin/streptomycin.

All the cell lines were grown at 37°C and supplemented with 5% CO_2_ in humidified incubators. The passage was carried out using 0.25% trypsin-EDTA (GIBCO) followed by centrifugation (1000 rpm, 4 min) and resuspension in a complete medium. Cell counting was performed using Countess automated cell counter with trypan blue exclusion (Thermo).

Cells were confirmed to be mycoplasma-negative (e-Myco plus Mycoplasma PCR Detection Kit, iNtRON Biotechnology).

WMs, a375, and RPEs cells were maintained in standard culture conditions (DMEM+10% FBS, vessel: Corning® Primaria™ 25cm2 Rectangular Canted Neck Cell Culture Flask with Vented Cap, PN: 353808). Passage was carried out using 0.25% trypsin-EDTA (GIBCO) followed by centrifugation (1000 rpm, 4 min) and resuspension in complete medium. Cell counting was performed using Countess automated cell counter with trypan blue exclusion (Thermo).

Prior to imaging/ proteomic analysis, cells were plated at day 0 in either 384-well PerkinElmer PhenoPlates (black, optically clear flat-bottom for imaging) or T175cm flasks for proteome analysis. For 384 wells the cell densities used per well were: T-47D (1200 cells), BT-474 (2400 cells), SKBR3 (2200 cells), 468 (1000 cells), 231 (800 cells), LM2 (800 cells), MCF10A (400 cells), and for the proteomics experiments they were scaled according to the surface area of the vessel used. Following three days of incubation in the above growth media, cells were either fixed in pre-warmed 4% formaldehyde (ThermoScientific) in PBS for 15 min at room temperature (image analysis) or collected in a pellet for proteomics analysis.

### B. Immunostaining

After fixation, cells were washed three times in PBS and then permeabilised in 0.2% Triton X-100/PBS solution for 15min at RT. Following three washes in PBS, cells were blocked for 1h in 2% bovine serum albumin (BSA) (Sigma)/PBS solution at RT. When using both mouse and rat primary antibodies in the same sample, sequential immunostaining was performed to avoid any antibody cross-reactions. Typically co-immunostaining with a mouse, rat and rabbit antibody was used. After the Block step, BSA was removed and the desired mouse primary antibody was added in Antibody solution (0.5%BSA/0.01% Triton X-100/PBS) at the indicated dilutions: YAP (G6) (Santa Cruz, 1:100), YAP/TAZ [67.3] (Santa Cruz, 1:1000). All the primary antibodies immunostainings were performed overnight at 4°C. Then cells were washed three times in PBS and incubated with a goat anti-mouse antibody 1:1000 in Antibody solution for 2h at RT. Cells were washed three times in PBS and incubated with a rat anti-tubulin alpha antibody (Bio Rad, 1:1000) and an anti-rabbit primary antibody when applied, for 2 hours at RT. The anti-rabbit primary antibodies were used at the indicated dilutions: TAZ (V386) (Cell Signalling, 1:200), Anti-PhosphoRB (Abcam, 1:1000). Then cells were washed three times in PBS, and incubated for 2h at room temperature with a goat anti-rat antibody and/or a goat anti-rabbit antibody or Alexa-488 phalloidin (Invitrogen) if needed. Finally, to stain nuclei, 5 mg/ml Hoescht (Invitrogen)/PBS solution was carried out for 15min at RT. 384-well plates were sealed for imaging with an Opera Cell:Explorer-automated spinning disk confocal microscope (PerkinElmer) or Opera Phenix (PerkinElmer) in the magnification indicated in the figure legends. At least twenty fields at random positions per well of a 384-well plate were imaged.

For the cell cycle experiments in MCF10A cells, samples were fixed in freshly prepared 4% PFA/PBS for 15 minutes. Cells were subsequently permeabilized with 0.25% Triton/PBS for 10 mins and blocked with 0.5% BSA/0.02% glycine/PBS for 30 minutes. Primary antibodies CCNA2 (Abcam, ab181591, 1:250) and YAP/TAZ (Santa Cruz, SC-101199, 1:250) were introduced via the same solution and left on for 1 hour at room temperature or overnight at 4 degrees. The plates were washed with PBS and the same was carried out for the secondary antibodies (Alexa fluor conjugated goat anti-mouse or anti-rabbit, 1:500) for 1 hour at room temp in PBS. Hoechst stain was added post-secondary (1:500) to stain DNA. Plates were imaged as above using the Opera Cell:Explorer with 20X objective lens (NA = 0.45).

### C. Image Acquisition and Feature Extraction

Image acquisition and cell segmentation was performed using Columbus high-content image analysis software or Harmony software. Nuclei were segmented using the Hoechst channel. Cell bodies were segmented using the tubulin channel. The perinuclear region was used to measure cytoplasmic antibody intensities. The cell-cell contact area (Neighbour fraction) was determined using an inbuilt Columbus algorithm ‘Cell Contact Area with Neighbors [%]’ expressed as the Percent of the object border that touches a neighbor object. The border objects were removed from the analysed cells considering only cells completely imaged. Mitotic cells were filtered using a combination of Hoechst intensity mean and Hoechst intensity maximum and excluded of all the analysis of this study. Geometric features measured include: the area of all subcellular regions; the length, width, and elongation (length/ width) of the cell and nucleus, cell and nuclear roundness and nucleus area/cytoplasm area.

#### Scaling Analysis

We conducted `k-means clustering` on the integrated Hoescht intensity of each cell line against nuclear area across `k = 1:8. k` was calculated using the elbow method and augmented with the additional constraint that cluster centroids should be separated by a factor of `2x`, where `x = 1:k`. This way, the first cluster approximates `2n G1`, the second `2n G2/4n G1` and so on. The YAP/TAZ – size relationship in each DNA-cluster was treated as a power law such that:

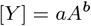

Where; [*Y*] represents YAP/TAZ concentration, ‘A’ cell area, and ‘a’ and ‘b’ are constants. The scaling factor, ‘b’ and ‘log(a)’ was extracted by conducting a linear fit on:

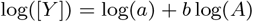

Which results from a simple manipulation. The factor by which ‘a’ increases across DNA groups is trivially retrieved by:

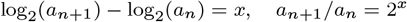

Examining the logarithmic derivative of [*Y*], we noticed that ‘b’ was not constant across the entire size range captured in our populations, although was over an 8-fold size difference about the mean. To avoid this complicating our analysis, we conducted the linear fits on the data within three standard deviations of the mean. Linear fitting, clustering, and data handling were conducted in the MATLAB R2019b (Mathworks) environment.

### E. Calculating Cell Area Distributions

Cell area distributions were derived from a simple adder system we published in a previous study (9). Briefly, we considered the probability of a cell advancing to the next cell cycle stage at any given time, `P`, and the cell’s growth rate, *β*, as proportional to the area at which it divided in the previous cycle:

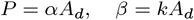

Where ‘*α*’ and ‘*k*’ are proportionality constants. From these two rules, it can be shown that in each cell cycle stage, the cell gains a random amount of mass drawn from a size-invariant exponential distribution centred on *k/α, R*(*A*), resulting in adder-like behaviour:

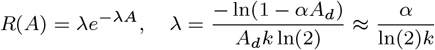

The division area distribution, *H*(*A*), results from the convolution of the birth distribution *B*(*A*) with *R*(*A*) ‘n’ times, where *n* is the number of cell cycle stages. Assuming symmetrical cell division, the subsequent *B*(*A*) (in the next cycle), is given as *H*(2*A*). This is then again convolved with *R*(*A*), and so on, resulting in a hypo-exponential distribution of general form:

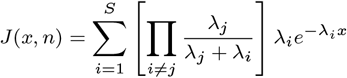

Substituting our values for a one-stage cycle system:

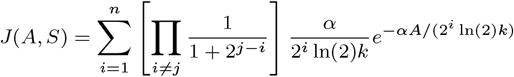

The n-stage distribution is obtained by first substituting *λ* for *nλ*:

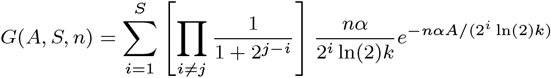

And convolving *G*(*A, S, n*) with itself ‘n’ times:

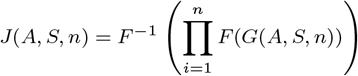

Where ‘F(c)’ denotes the Fourier transform of ‘c’ and ′*F* ^(^ 1)(*c*)′ the inverse. Here we have leveraged the convolution theorem to express the convolution as a multiplication in Fourier space. For calculation, the initial cell area distributions are considered a delta function centred on ln(2)*k/α* (the mean of *R*(*A*)). Every generation, the area distribution is convolved with the mass-gain distribution, *R*(*A*), ‘n’ times, computed by performing an inverse Fourier transform on the product of the two distributions’ respective Fourier transforms. This produces the division area distribution, *Ad*(*A*), which must be transformed to *Ad*(2*A*) to capture the effects of cell division. We perform this by setting *Ab*(*Ax*) = *Ad*(*Ai*) + *Ad*(*Ai* + 1), where ‘i’ = xn-xn-1 for all *x*, where *Ab* denotes the birth size distribution. This is then convolved with the gain distribution as before to generate the next division distribution and so on until a desired number of generations has been reached. For each, we calculated the Kullbeck-Liebler divergence between the experimental and simulated data to assess model error. For discrete probability distributions defined on the same probability space, *X*, the Kullback–Leibler divergence from *P* to *Q* is:

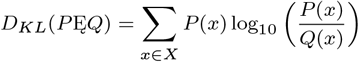

Model fitting was conducted within the commercial `MATLAB R2019b (Math Works)` software’s machine learning toolbox.

### F. Calculating YAP/TAZ Concentration Distributions

We computed the dilution factor distribution for an initial size condition, constrained such that the cell begins the simulation at its ‘expected size’ within our 2-stage adder proliferation model framework:

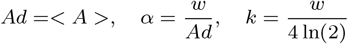

Where ‘w’ is an arbitrary constant. These parameters are used to compute an initial dilution factor distribution:

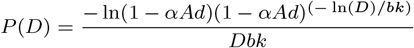

From which we compute the product distribution (assuming identical cell cycle stages):

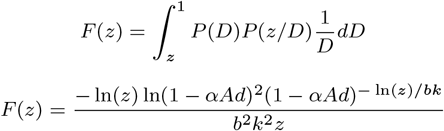

Where we assume that the cell grows approximately equal amounts in G1 vs G2. We then draw from this distribution, obtaining a dilution factor ‘d’, by passing a uniformly distributed random number through the inverse of *F* (*z*), defined as:

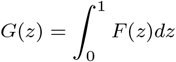

We multiply the initial YAP concentration by ‘K*d’ to generate the starting YAP concentration for the next cycle. ‘K’ is taken from the size-K curve experimentally measured. From this, we also calculated the size change needed to generate ‘d’ as:

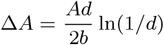

From which we trivially update *Ad*:

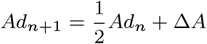

This now facilitates re-calculation of *P* (*D*), *F* (*z*), etc. for the next generation, and the cycle is repeated for 10 generations to generate one sample YAP concentration for the cell line. The process is repeated 1000 times to generate a YAP/TAZ concentration distribution for each line, which is compared to the experimentally determined YAP/TAZ intensity distributions via the KBL divergence.

### G. Calculation of the Coefficient of Variation of *F* (*z*)

To calculate the coefficient of variation of *F* (*z*), we first derived the mean:

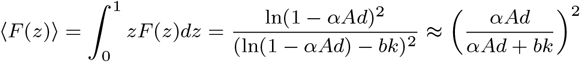

Where we have expanded about *α* = 0. The variance is given as:

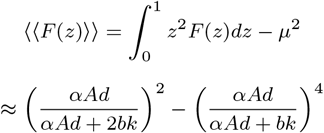

Where we have made the same simplification. From these, we obtain the coefficient of variation as:

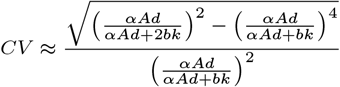

### H. PLSR and Hit Detection

Regression analyses were conducted with the MATLAB (MathWorks) environment using the plsregress function from the machine learning toolbox. Partial least squares regression was selected as the method to help mitigate the influence of co-linearity in the predictor dataset. Model components were selected through 10-fold cross-validation using the elbow method on the mean square error as a function of component number.

For the RNAi screening data, all 114 shape features were mean-centered prior to model construction. Models were built from control data and applied to the combined knockdown-control state. Fit quality was assessed through the r-squared metric. Linear models (predicted vs observed) were visualized through the ‘dscatter’ function. Z scores were calculated for the difference between observed and predicted YAP/TAZ ratios/abundances and the increase/decrease from the mean YAP/TAZ ratio/abundances. Knockdown ‘Hits’, those which decoupled, were selected from this analysis as knockdown states achieving an average *Z score >* 2 (two standard deviations from the mean).

For the proteomic analysis, phosphopeptide abundances were adjusted to reflect ‘excess’ phosphorylation given the total expression of the peptide. To do this, a regression model was constructed for each gene relating phosphopeptide and peptide abundance. The adjusted phosphopeptide abundance was taken as the phosphopeptides’ deviation from this regression model.

#### 1. Feature Importance to PLSR Models

The influence a feature has on a model was estimated through ‘Variable importance to projection’ (VIP) scores calculated as:

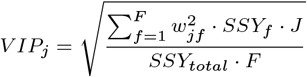

Where *w*_*jf*_ is the weight value for the *j* variable and *f* component, *SSY*_*f*_ is the sum of squares of explained variance for the *f* th component, *J* number of *X* variables, and *SSY*_*total*_ is the total sum of squares explained of the dependent variable, and *F* is the total number of components. Features with a VIP score greater than 1 were taken as major drivers of the models.

### J. Linear Classifier

For the cell cycle experiments in MCF10A cells (engineered to express endogenous mRuby-tagged PCNA) we used a manually trained linear classifier. Cell cycle classification was performed using Columbus (PerkinElmer). We used a combination of thresholding and linear classifiers based on nuclear morphology and DNA, CCNA2, and PCNA intensity and texture features. Classification was performed sequentially by manual annotation to divide and further subdivide cell cycle stages. First, nuclei and cell bodies were segmented using the DNA and YAP/TAZ channels and cells touching the border were removed. Then mitotic nuclei were distinguished from interphase nuclei based primarily on DNA, PCNA and morphology features using a manually trained linear classifier (most relevant features: nucleus DNA texture Bright/Edge/Ridge, PCNA intensity, nucleus area/roundness/width, nucleus DNA intensity). Interphase nuclei were thresholded based on mean nuclear PCNA intensity, with PCNA-nuclei classed as G0. PCNA+ nuclei were divided into CCNA2+ and CCNA2-subpopulations based on mean nuclear CCNA2 intensity and PCNA+/CCNA2-cells were classed as G1. PCNA+/CCNA2+ cells with low mean CCNA2 intensity (first quartile) were classed as Early S-phase. The remaining cells were finally divided into S and G2 classes using a manually trained linear classifier. During S-phase, PCNA goes from being uniformly distributed in the nucleus to having a progressively more punctate or spotty appearance as DNA replication proceeds. The PCNA texture linear classifier was manually trained on PCNA+/CCNA2high cells (most relevant features: PCNA texture Edge/Saddle/Ridge/Haralick Homogeneity/CV, mean nuclear CCNA2 intensity, mean perinuclear ring region CCNA2 intensity). “Spotty” nuclei classed as S-phase and “smooth” nuclei classed as G2. YAP/TAZ intensity features were not included in the spotty/smooth linear classifier. Integrated DNA intensity (i.e. total amount of DNA) was not included in the spotty/smooth linear classifier but was used post-hoc to verify S versus G2 classification.

### K. Cell preparation and proteomics analysis of the Breast cell lines

Cells were plated at day 0 as stated above and collected 72h later by trypsinization. After resuspension in growth media and centrifugation, media was removed and 1mL of cold PBS was added. Then one million of viable cells per cell line (by duplicate) was transferred to low binding tubes and washed 2 x with cold PBS to a final pellet that was flash frozen with 70% ethanol and dry ice. Cell pellets were lysed in 1% sodium deoxycholate (SDC), 100 mM triethylammonium bicarbonate (TEAB), 10% isopropanol, 50 mM NaCl buffer freshly supplemented with Halt protease and phosphatase inhibitor cocktail (100X) (Thermo, 78442), 5 mM tris-2-carboxyethyl phosphine (TCEP), 10 mM iodoacetamide (IAA) and Universal Nuclease (Pierce) followed by bath sonication for 5 min and incubation at room temperature for 45 min. Protein concentration was measured with the Quick Start Bradford protein assay. Aliquots of 60 g of total protein were digested overnight with trypsin (Pierce, ratio 1:20) and labelled with the TMTpro multiplex reagents (Thermo) according to manufacturer’s instructions. The peptide mixture was fractionated with high pH Reversed-Phase (RP) chromatography using the XBridge C18 column (2.1 x 150 mm, 3.5 m, Waters) on an UltiMate 3000 HPLC system over a 1% gradient in 35 min. Mobile phase A was 0.1% (v/v) ammonium hydroxide and mobile phase B was 0.1% ammonium hydroxide (v/v) in acetonitrile. Phosphopeptide enrichment was performed with the High-Select™ Fe-NTA Phosphopeptide Enrichment Kit (Thermo) using a modified protocol in a well plate array format. A volume of 50 L resin/buffer was transferred on top of 10 L filter tips that were fitted on a 96-well plate using a suitable tip rack. The resin was washed three times with 40 L wash/binding solution and centrifugation at 500 g for 1 min. Peptides were reconstituted in 30 L wash/binding solution and were loaded onto the tip-columns with the resin. After 30 min, the flow-through (FT) from three washes with wash/binding solution were collected in a clean 96-well plate with centrifugation at 500 g for 1 min each time. Phosphopeptides were eluted twice with 40 L elution buffer in a clean 96-well plate with centrifugation at 500 g for 1 min, transferred in glass vials (Waters, P/N 186005669CV) and SpeedVac dried. Both the flow-through solutions and IMAC eluents were subjected to LC-MS analysis for bulk proteome and phosphoproteome analysis respectively. LC-MS analysis was performed on an UltiMate 3000 system coupled with the Orbitrap Fusion Lumos Mass Spectrometer (Thermo) using an Acclaim PepMap, 75m x 50cm C18 capillary column over a 95 min (FT) or 65 min (IMAC elution) gradient. MS spectra were collected with mass resolution of 120k and precursors were targeted for HCD fragmentation in the top speed mode with collision energy 36% and IT 54 ms (FT) or 100 ms (IMAC elution) at 30k Orbitrap resolution. Targeted precursors were dynamically excluded from further activation for 45 or 30 seconds. The Sequest HT engine in Proteome Discoverer 2.4 (Thermo) was used to search the raw mass spectra against reviewed UniProt human proteins. The precursor mass tolerance was set at 20 ppm and the fragment ion mass tolerance was 0.02 Da. TMTpro at N-terminus/K and Carbamidomethyl at C were defined as static modifications. Dynamic modifications were oxidation of M and deamidation of N/Q as well as phosphorylation of S/T/Y for the phosphoproteome analysis. Peptide confidence was estimated with the Percolator node and peptide FDR was set at 0.01. Only unique peptides were used for quantification, considering protein groups for peptide uniqueness. Peptides with average reporter signal-to-noise greater than 3 were used for protein quantification.

### L. Gene Set Enrichment Analysis

GSEA was conducted using the ‘WebGestalt’ web application on our ranked list of peptides (VIP-Score defined the rank) (76). We used the ‘pathway’ and ‘noRedundent Biological process’ enrichment categories to identify enriched themes/pathways in the high and low scoring peptides. Parameters used: Minimum IDs per category =5, max = 10000, permutations = 1000. Enrichments with a false discovery rate *<* 0.05 were taken as ‘hit’ themes and/or pathways.

### M. Drug treatments

231 and LM2 were plated in 384 wells and treated with 10uM of Binimetinib or DMSO. 24h after cells were fixed in pre-warmed 4% formaldehyde (ThermoScientific) in PBS for 15 min at room temperature. For the Palbociclib experiments in LM2, the drug was used at 0.5 uM for 24h hours prior to formaldehyde fixation, immunofluorescence, and image analysis.

For the experiments in melanoma cells and RPEs 0.33uM Palbociclib and 0.25uM Binimetinib were added to the cell cultures and incubated for a duration of 72 hours. Following the treatment, the cells were fixed in 4% paraformaldehyde (PFA) for 15 minutes at room temperature. Primary antibody staining was performed using a dilution of 1:1000 for YAP/TAZ and 1:1000 for pRB. Secondary antibody staining was conducted using a dilution of 1:500. All antibody stains were incubated overnight at 4 degrees Celsius.

### N. Data Availability

Image datasets for the cell lines used for morphological profiling are available from: DRYAD: http://dx.doi.org/10.5061/dryad.tc5g4.

Image Data Repository (http://idr-demo.openmicroscopy.org/about, accession number S-BSMS6)

Biostudies database (https://www.ebi.ac.uk/biostudies/studies/S-BSMS6).

## 3. Author Contributions

I.Jones analysed the data, developed the computational models and with C.Bakal wrote the manuscript. J.Sero, M.Arias-Garcia and M.Beykou cultured the breast cells and generated the imaging datasets. J.Sero and M.Arias-Garcia carried out the image analysis. P.Pascual-Vargas conducted the RNAi screens and performed the image analysis and M.Arias-Garcia optimised the screening and staining methodology. M.Arias-Garcia and L.Dent performed the drug experiments and their image analysis in breast and melanoma cells respectively. T.Roumeliotis, M.Beykou and M.Arias-Garcia, with J.Choudhary, generated the proteomic datasets. C.Bakal conceived and designed the research.

## Supplemental Information

A guide to the datasets used in this study:

- YAP/TAZ concentration decreases with increasing cell size: Sero et al, mol sys bio, 2015
- YAP/TAZ abundance, concentration, and N/C ratio scaling are insensitive to cell crowding: Sero et al, mol sys bio, 2015
- YAP/TAZ concentration, but not scaling, is sensitive to DNA-content and cell cycle progression: Sero et al, mol sys bio, 2015 / Sero novel to this study
- A constant nuclear concentration of YAP/TAZ is maintained across cell sizes despite whole cell dilution: Sero et al, mol sys bio, 2015
- the nuclear YAP/TAZ concentration is associated with altered RAS and nuclear transport signalling processes: M.Arias-Garcia, M.Beykou, novel to this study
- YAP/TAZ dilution behavior is conserved across melanoma and RPE cells: L.Dent, T. Pal Chaudhuri, novel to this study
- 231/LM2 after the MEKi and LM2 after the CDK4/6 inhibitor. M.Arias-Garcia, novel to this study
- Integration of YAP/TAZ size-scaling and stochastic cell division fosters YAP/TAZ heterogeneity: Sero et al, mol sys bio, 2015
- YAP/TAZ scaling behavior drives increasing nuclear/cytoplasmic ratios following gene depletion: P.Pascual-Vargas, Scientific data, 2017

### A. YAP/TAZ abundance, concentration and N/C ratio scaling are insensitive to cell crowding

Cell size and YAP/TAZ activation have previously been associated with cell crowding, cell-cell adhesion, and contact inhibition (34). In our cell lines, cell area negatively correlated with neighbour fraction (NF), raising the possibility that the observed YAP/TAZ sub-scaling was driven by the NF (Supp.fig. 3). To investigate this, we first quantified the extent to which YAP/TAZ scaled with NF, noticing a strong positive association at high NF’s (0.4 -1), in accordance with the area correlation. However, when clustering the cells on NF, and conducting a scaling analysis between YAP/TAZ concentration and cell area within each NF group, we observed no change in the YAP/TAZ concentration per size or ‘b’ across clusters (Supp.fig. 3). We conducted the same analysis for YAP/TAZ abundance and nuclear/cytoplasmic ratio, noting the same effect, with the exception that MCF10A’s YAP/TAZ N/C ratio exhibited a NF sensitivity (Supp.fig. 3) as is consistent with previous works (22). From these data, we concluded it is the NF’s influence on cell area that is driving the NF-YAP scaling relationship rather than any direct effect of NF on YAP/TAZ in these lines.

### B. YAP/TAZ scaling behavior is consistent with a dilution-synthesis-dilution scheme

Theoretically, we assume a two-stage system. Through stage 1 (G1), the cell area increases by a factor of *x*, and in the subsequent stage (G2), by a factor 2*/x* (such that the cell doubles in size across a cell cycle). The expected dilution factor, *F*_*G*1_, of YAP/TAZ through S1 as:

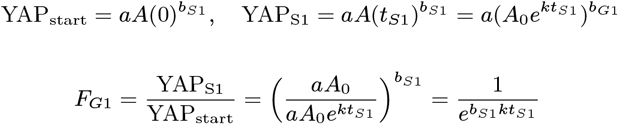

Although a suitable value of *k*, between stage synthesis, can mitigate any value of scaling factor, the DSD system does not behave identically for any valid parameter combination. This can be seen by integrating the DSD model with a simple adder system (9). Within this framework, the cell is considered to have a probability to advance cell cycle stage proportional to its division size:

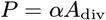

Leading to a cumulative distribution function for the proliferation time distribution given as:

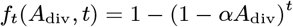

Expressing the proliferation time distribution in terms of the dilution factor *D*:

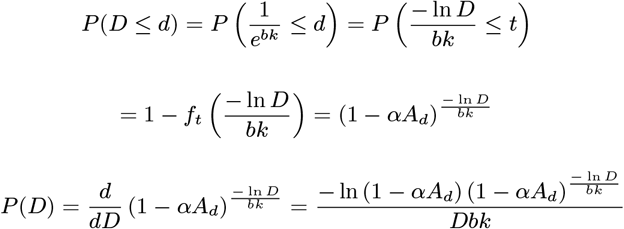

We obtain the probability distribution of dilution factors. As two dilution events happen in sequence, we are interested in the product distribution of *P* (*D*) and a second dilution event governed by a distribution *G*(*D*) given as:

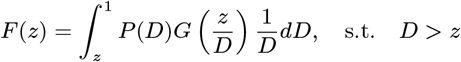

### C. YAP/TAZ scaling rate is a function of cell size

While we observed clear sub-scaling behavior when investigating the average scaling factor of YAP/TAZ with cell size, we were interested in how this extended to the case where the scaling factor *b* is treated as a continuous function of cell size, *b*(*A*). We extracted *b*(*A*) by taking the logarithmic derivative of YAP/TAZ concentration with respect to the cell area:

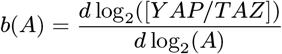

Strikingly, no cell line exhibited a constant scaling factor. Scaling factors (*b*) tended to be lower in small cells and steadily decreased with increasing cell size (within a cell line); however, in several lines, this relationship would reverse at larger sizes, with further increases in area leading to weaker sub-scaling. This extended to both the G1 and G2 populations. Remarkably, in a subset of our lines (e.g., HCC1954, MDA-MB-231), YAP/TAZ concentration even positively scaled with cell size at small sizes. Although, this effect was limited to G1 cells (Supp.fig. 10) implying the existence of an area-dependent scaling trigger.

To understand how significant size-variable scaling is to the functioning of the cell, we calculated the range of scaling factors that occur over the most common size “bands,” which we define here as the mean G1/G2 sizes +/- 1 std deviation. Calculating this for the G1 cells, we observed that 5/9 of our cell lines exhibited only a modest variation in the scaling factor within this size constraint (stddev = 0.03-0.08). However, the remainder showed far more extreme variations (stddev = 0.14-0.17) leading to an approximate 2-fold change in scaling factor *b* across the size range (Supp.fig. 10). The mean scaling rate was approximately constant across cell lines (−0.4 - -0.5). In the G2 group, the variation in scaling rate increased (std = 0.10-0.18) for all lines, and the mean scaling rate significantly decreased for only a subset (Supp.fig. 10). Thus, the YAP/TAZ scaling factor varies within relevant size ranges and should not be considered a constant.

We also investigated how the fold difference in YAP/TAZ concentration between ‘G1’ and ‘G2’ cells, *K*, varied with cell size. In most (7/9) of our cell lines, *K* was found to decrease with increasing cell size; at small sizes (mean size – 2std deviations) taking values between 1.3-1.6, and at larger sizes (mean size + 2std deviations), 1-1.3 (Supp.fig. 10).

Together, these data show that the scaling of YAP/TAZ with cell size is not static but changes dynamically with cell size within relevant size ranges in both G1 and G2 cells. This size-dependence extended to the fold-change across DNA-contents, *K*, suggesting that the size at which a cell passes the G1/S checkpoint informs the scaling and production of YAP/TAZ across subsequent cell cycle stages and potentially, generations.

### D. An increased nuclear YAP/TAZ concentration correlates with a reduction in the YAP/TAZ scaling factor

We noticed that lines exhibiting super-scaling behavior in their nuclear YAP/TAZ concentrations at small cell sizes were those with the most variable scaling factors. To investigate this, we began by assuming a linear relationship between the scaling factor *b* and the logarithm of the nuclear YAP/TAZ concentration:

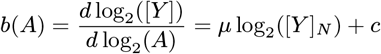

Where *μ* and *c* are constants to be determined through fitting to the experimentally determined *b*(*A*) and *Y*_*n*_. As *b*(*A*) and *Y*_*n*_ vary with size only at very small sizes, when fitting to this equation, we considered only cells meeting the criterion *A*_*c*_ < ⟨*A*⟩ − *σ*, where ⟨*A*⟩ is the mean cell area and *σ* is the standard deviation. This avoided the greater density of points at a constant *b* and *Y*_*n*_ from skewing the calculation. We used equ.S9 to calculate the expected scaling rate at each cell area and found excellent agreement with that measured (Supp.fig. 12). Importantly, while an increase in nuclear YAP/TAZ leads to a decrease in scaling factor, a decrease in YAP/TAZ leads to an increase in scaling factor (see SUM149 and MDAMB157 Supp.fig. 12). This suggests that the effect is not driven by a correlate of YAP/TAZ import but by a correlate of the nuclear concentration itself.

This effect did not extend across lines; higher nuclear YAP/TAZ concentrations did not lead to lower scaling factors. Instead, scaling factors were more associated with higher YAP/TAZ ratios. The direction of causality is unclear, but it is plausible that higher scaling factors induce greater ratios by virtue of reducing the cytoplasmic YAP/TAZ concentration (Supp.fig. 12).

Together, this suggests that the correlation between the nuclear YAP/TAZ concentration and scaling factor may emerge from a co-dependence on an unseen cryptic variable rather than any direct effect of the nuclear YAP/TAZ concentration. Indeed, as Palbociclib treatment increases nuclear YAP/TAZ whilst delaying whole cell dilution to larger sizes, it is likely that the correlation between the nuclear YAP/TAZ concentration and the scaling factor emerges from nuclear translocation occurring before division commitment and that same commitment triggering whole-cell dilution. However, we cannot yet exclude the possibility of a negative feedback mechanism.

**S.Figure 1:**
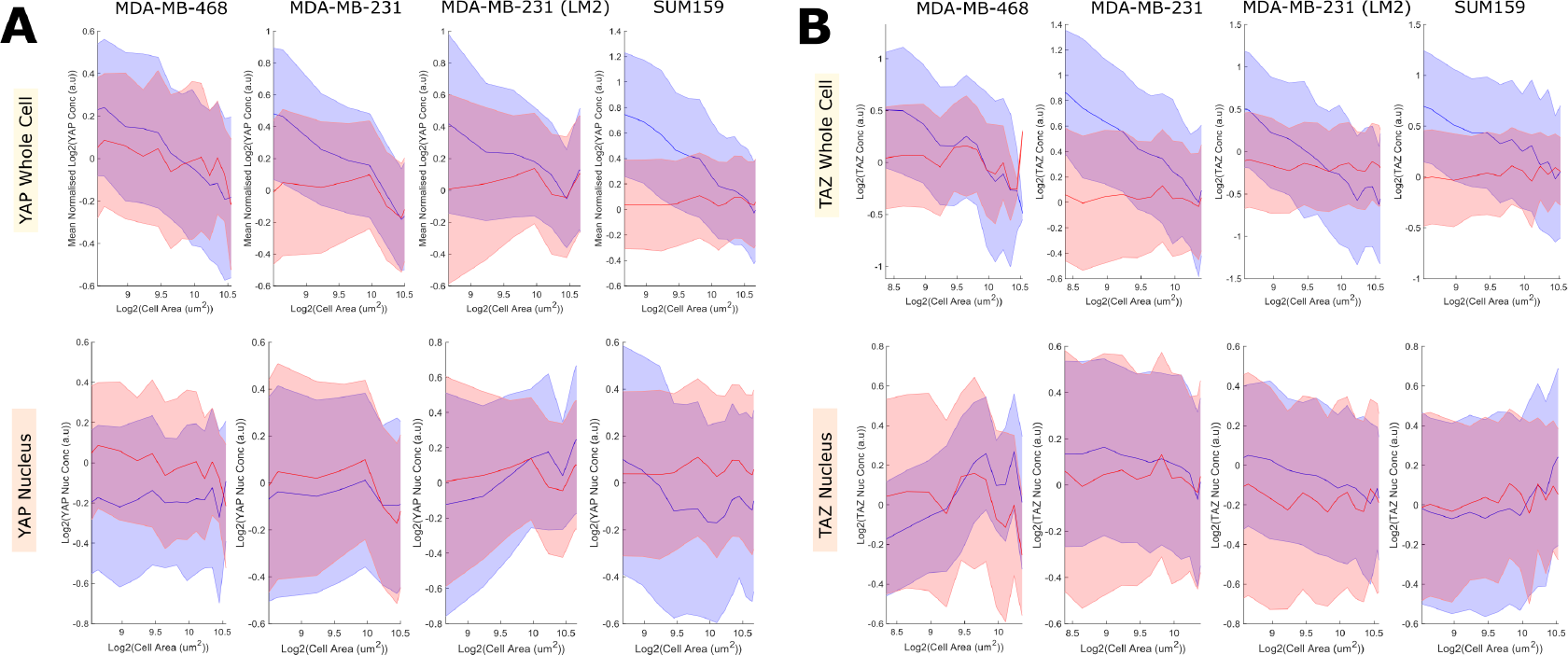
Deconvolved YAP and TAZ size-scaling A) Log-log plots relating YAP (blue) and Actin (red) abundance to single-cell area. Assuming a power-law relationship between protein concentration and cell size, the gradient represents the power to which concentration scales with area and the y-intercept determines the maximum concentration. The shaded area denotes one standard deviation of the cell size distribution about that size bin. B) Log-log plots relating TAZ (blue) and Actin (red) abundance to single-cell area. Assuming a power-law relationship between protein concentration and cell size, the gradient represents the power to which concentration scales with area and the y-intercept determines the maximum concentration. The shaded area denotes one standard deviation of the cell size distribution about that size bin.

**S.Figure 2:**
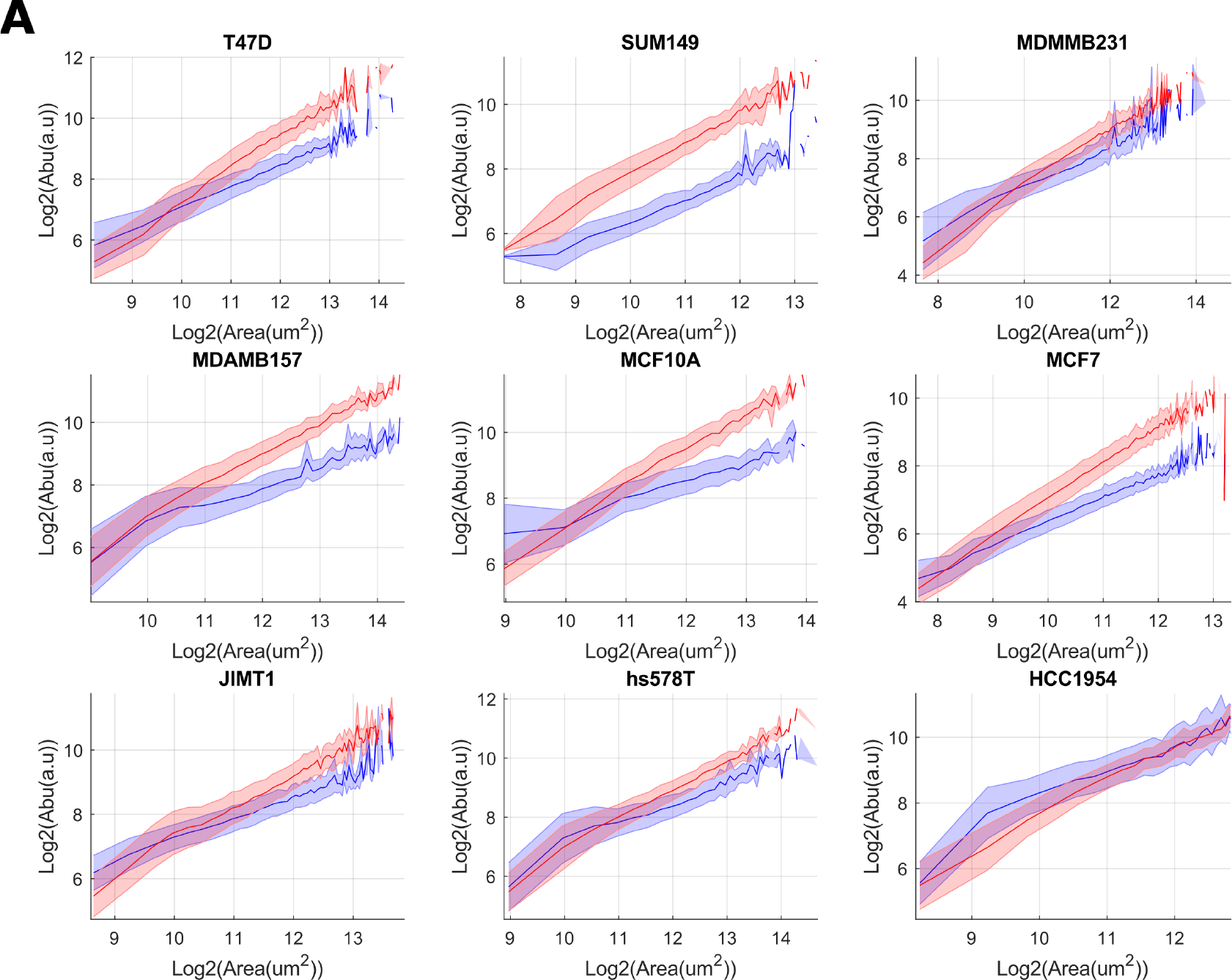
YAP/TAZ and actin abundance-size scaling. A) Log-log plots relating YAP/TAZ (blue) and Actin (red) abundance to single-cell area. Assuming a power-law relationship between protein concentration and cell size, the gradient represents the power to which concentration scales with area and the y-intercept determines the maximum concentration. The shaded area denotes one standard deviation of the cell size distribution about that size bin.

**S.Figure 3:**
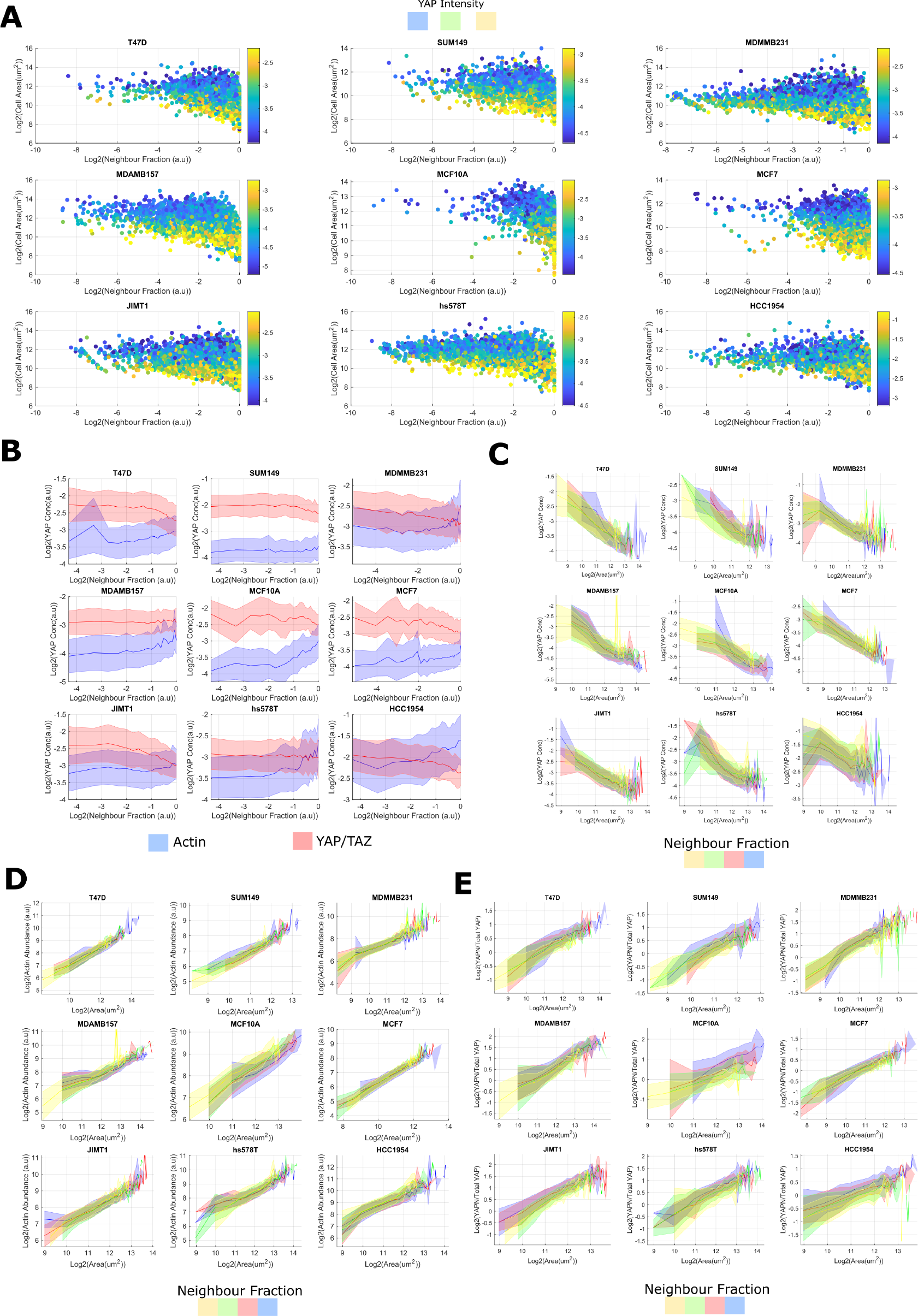
The effect of neighbour fraction on YAP/TAZ scaling A) Log-log plots relating single cell neighbour fraction and cell area. Colour is proportional to the mean YAP intensity (concentration). Area and NF negatively correlate at high (close to 1) neighbour fractions. B) Log-log plots relating YAP/TAZ (blue) and Actin (red) concentration to single-cell neighbour fraction. Assuming a power-law relationship between protein concentration and cell size, the gradient represents the power to which concentration scales with NF and the y-intercept determines the initial concentration at NF = 0. The shaded area denotes one standard deviation of the cell size distribution about that size bin. C) Log-log plots relating YAP/TAZ concentration and single cell area across lines and each NF bin (as determined by kmeans clustering on the neighbour fraction). Blue represents the lowest NF, then red, green and yellow, the most. The shaded area denotes one standard deviation of the cell size distribution about that size bin. D/E) As in ‘C’ but relating to YAP abundance (D) or N/C ratio (E).

**S.Figure 4:**
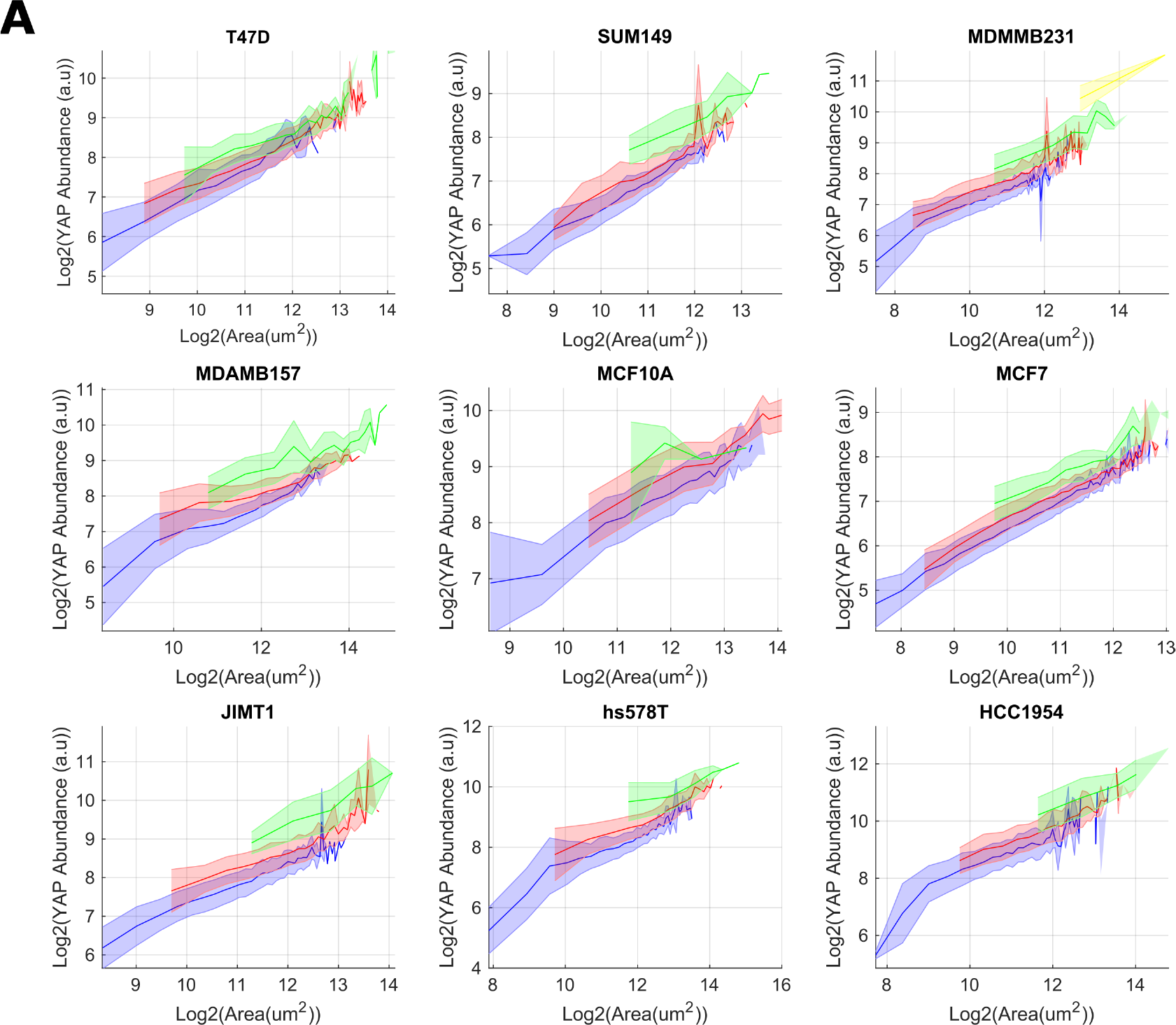
DNA sensitivity of YAP/TAZ abundance-size scaling A) Log-log plots relating YAP/TAZ abundance and single cell area across lines and each DNA content bin (as determined by kmeans clustering on the integrated Hoechst intensity and nuclear area). Blue represents the lowest DNA content, then red, green and yellow, the most. The shaded area denotes one standard deviation of the cell size distribution about that size bin.

**S.Figure 5:**
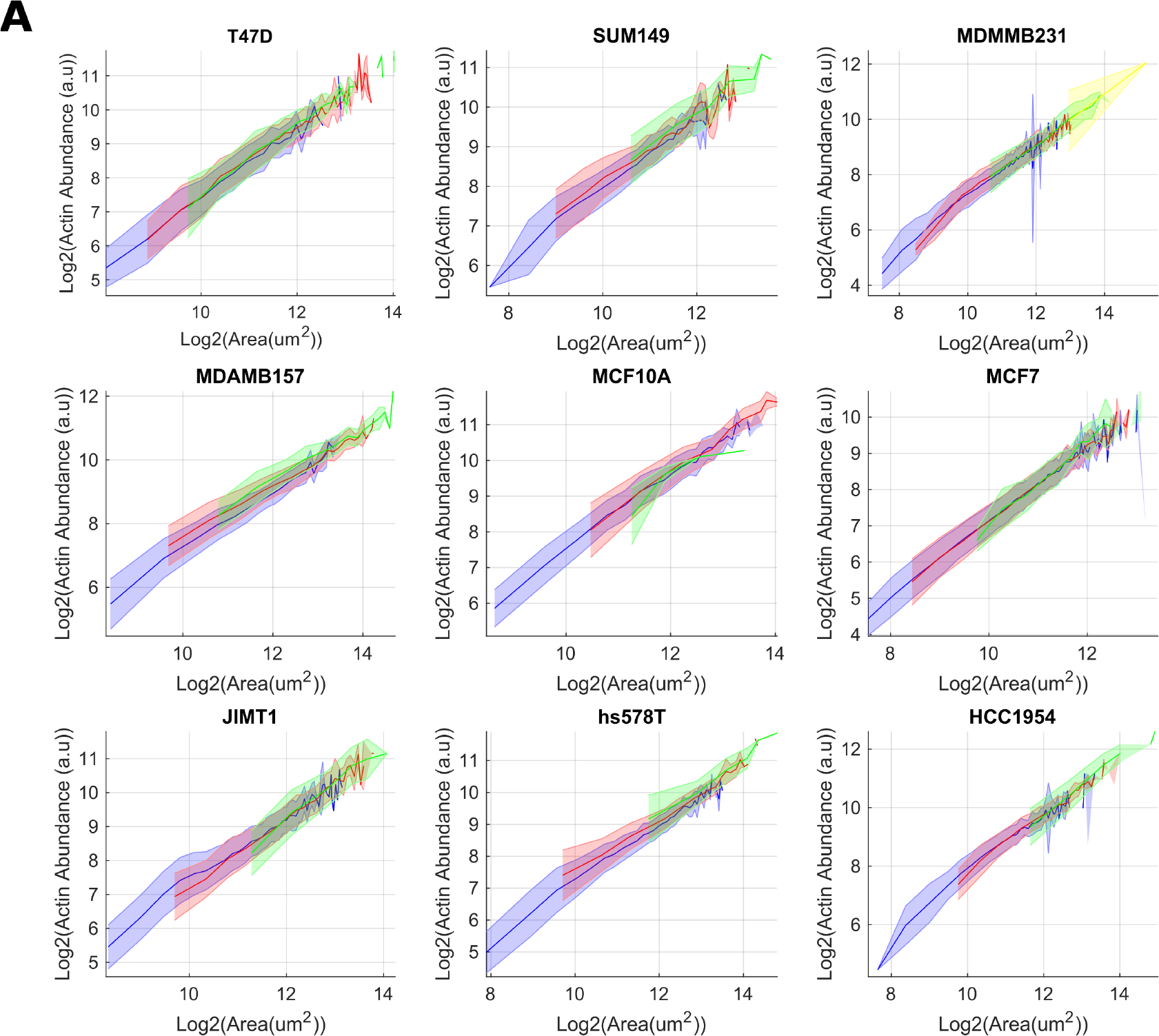
A) Log-log plots relating Actin abundance and single cell area across lines and each DNA content bin (as determined by kmeans clustering on the integrated Hoechst intensity and nuclear area). Blue represents the lowest DNA content, then red, green and yellow, the most. The shaded area denotes one standard deviation of the cell size distribution about that size bin.

**S.Figure 6:**
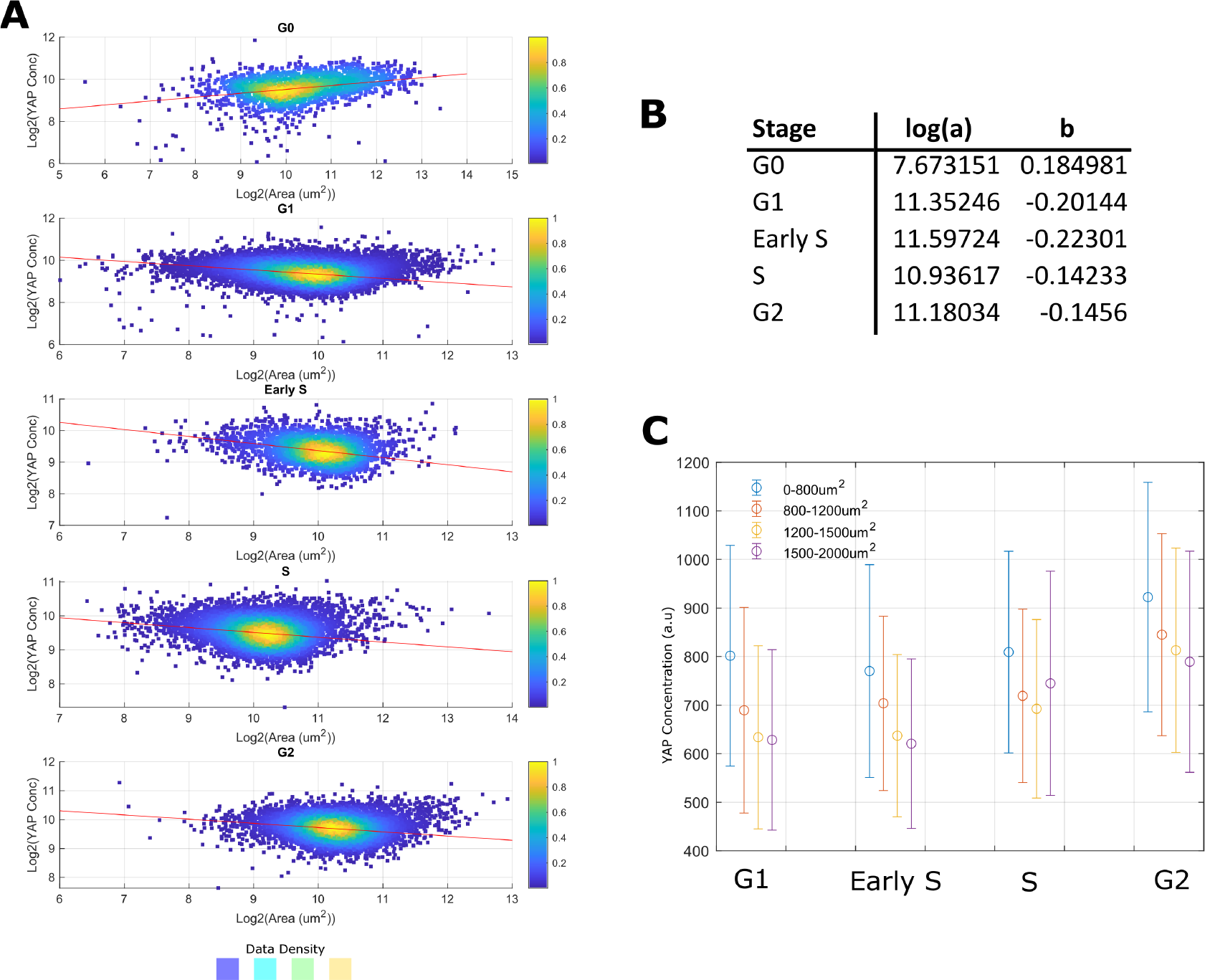
YAP/TAZ scaling is independent of cell cycle phase A) Log-log plots relating single cell area and YAP/TAZ concentration across cell cycle stages. Stages are defined by a linear classifier trained on PCNA and CCNA1 intensity data. Colour is proportional to the density of the data. The red line is a linear fit of the data. B) Scaling parameters for each cell cycle stage; log(a) is the y-intercept and ‘b’ is the gradient. C) Mean YAP/TAZ concentration across cell cycle stages where cells are ‘binned’ by size (indicted by colour).

**S.Figure 7:**
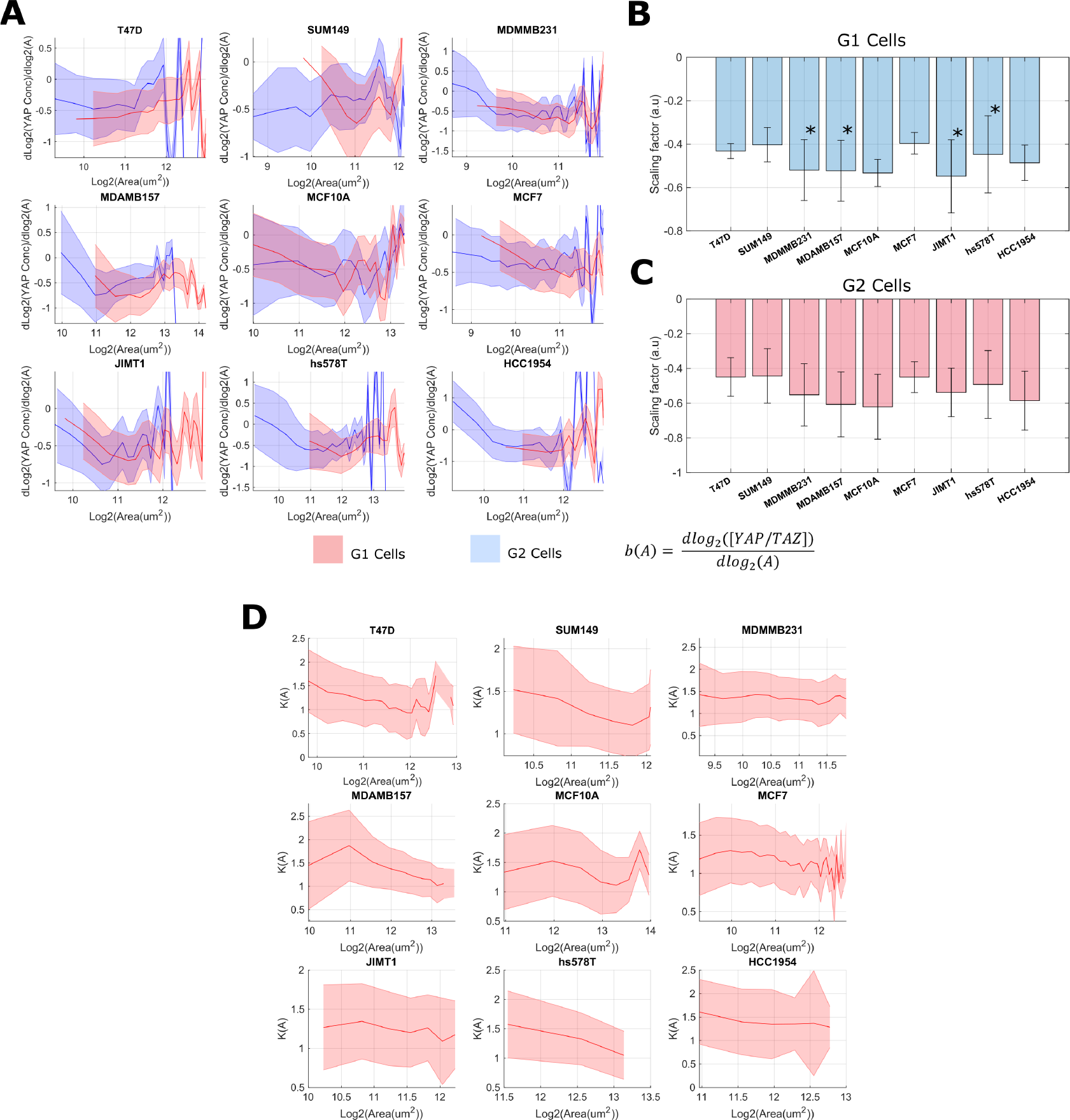
YAP/TAZ scaling rate is a function of cell size: A) The relationship between Log(cell area) and the logarithmic derivative of YAP concentration with respect to area. The blue line represents G1 cells, whilst the red represents G2 cells. G1 and G2 groups were determined through kmeans clustering on the integrated Hoechst intensity. Shaded regions represent one standard deviation of all values within the local size ‘bin’. B) Average scaling factors within one standard deviation of the G1 area distribution mean. Error bars represent one standard deviation of the scaling factor distribution in that size range. Cell lines marked with an asterisk are those that showed the highest variance in scaling factor within a ‘typical’ size range, in each case, these cell lines show a larger than average scaling factor at small sizes. C) Average scaling factors within one standard deviation of the G2 area distribution mean. Error bars represent one standard deviation of the scaling factor distribution in that size range. D) The fold-change between G2 and G1 YAP/TAZ as a function of cell size. The shaded region represents one standard deviation from the mean.

**S.Figure 8:**
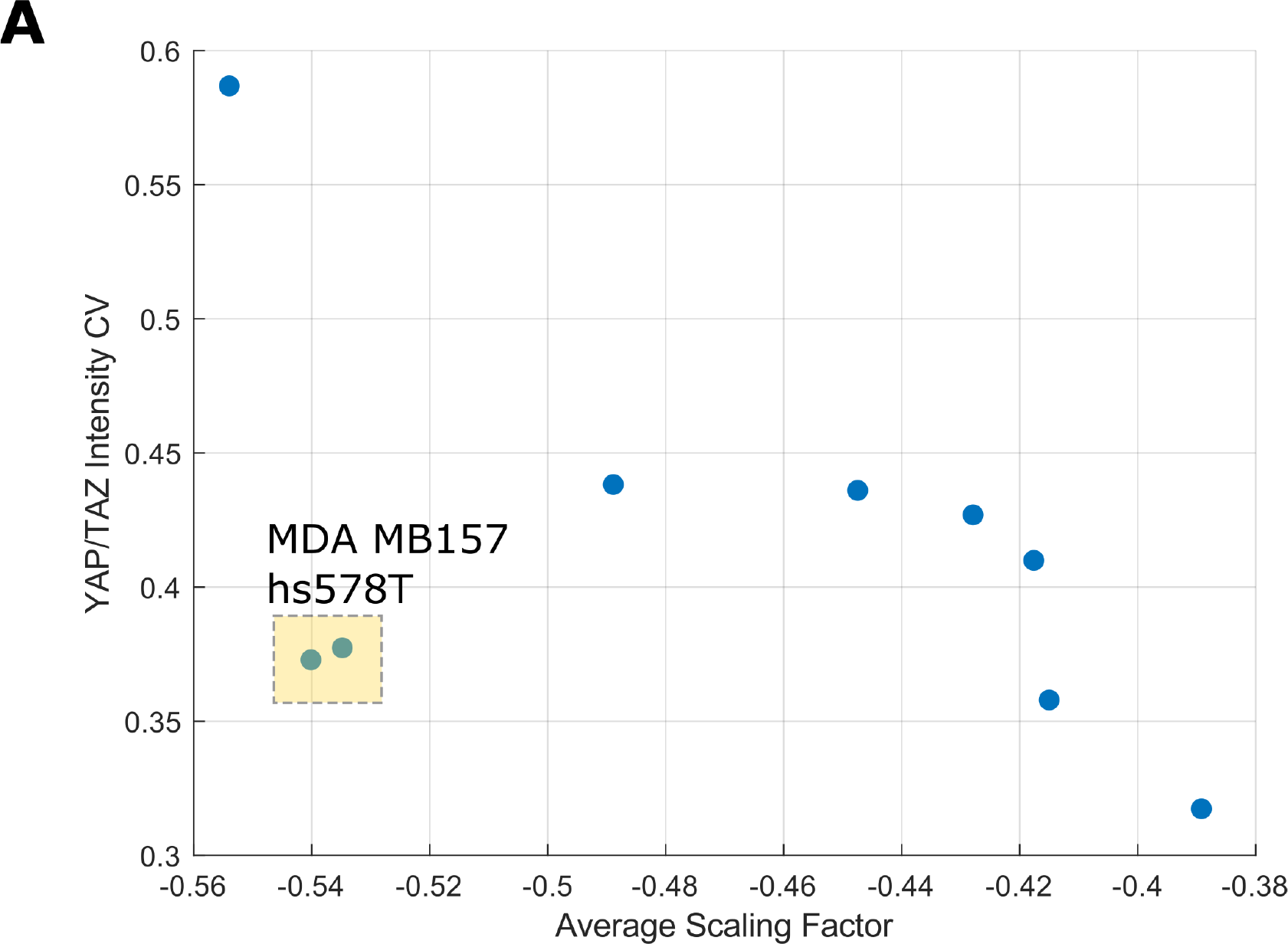
YAP/TAZ intensity distribution variance and the scaling factor. The relationship between the coefficient of variation of the YAP/TAZ intensity distributions and the average scaling factor across G1 and G2 populations.

**S.Figure 9:**
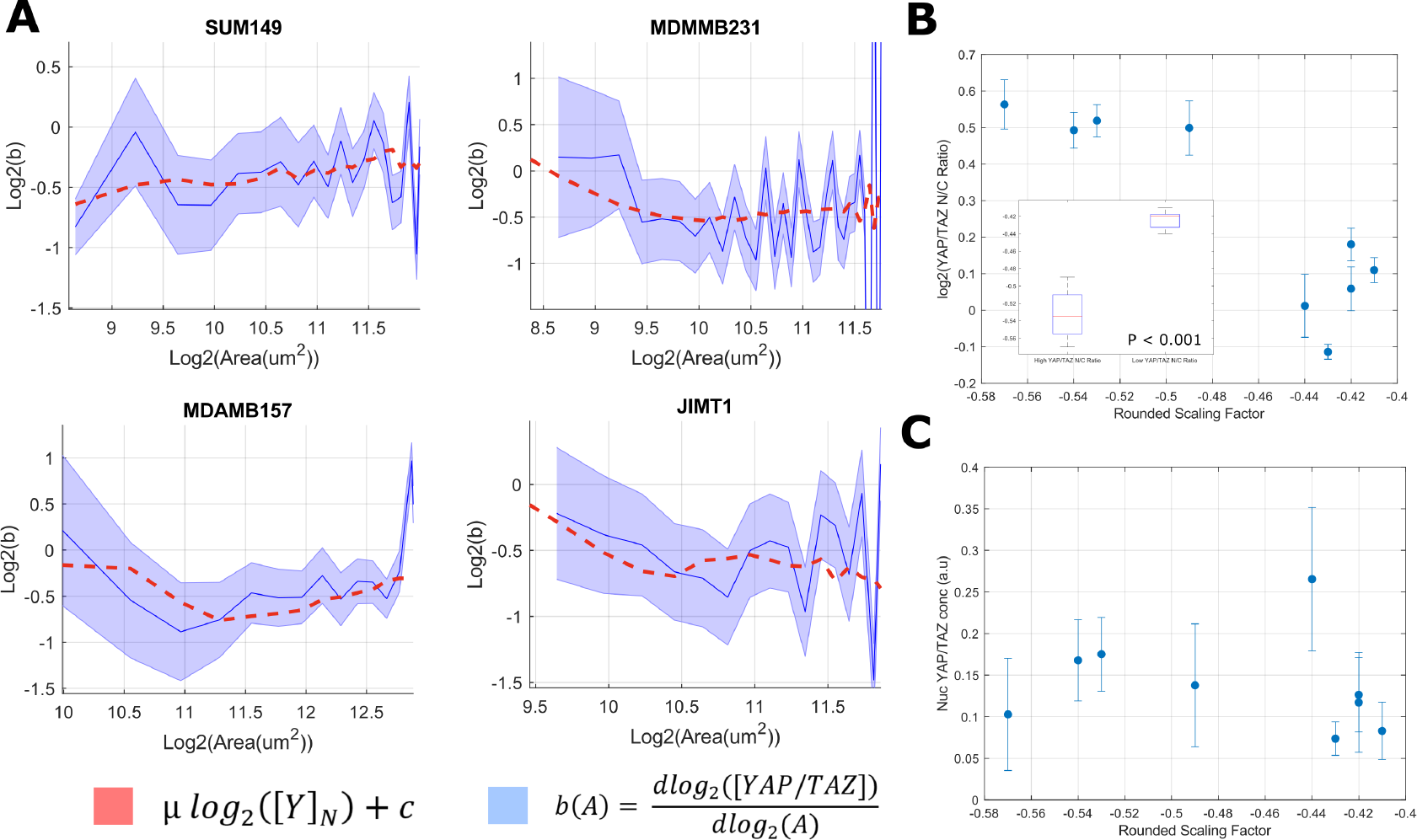
Nuclear translocation of YAP/TAZ correlates with a reduction in YAP/TAZ scaling factor. A) Relationship between total YAP/TAZ scaling factor, ‘b’, and cell area (blue) in the G1 cell population (as determined by kmeans clustering on the integrated Hoechst intensity). Cell lines shown are those which exhibited the greatest area sensitivity in their scaling factors. The error margin corresponds to 1 standard deviation in that size bin. The red line is an estimate of ‘b’ assuming a linear relationship between ‘b’ and log(Yn), where Yn is the nuclear YAP/TAZ concentration. B) YAP/TAZ nuc/cyto ratio against the average cytoplasmic YAP/TAZ scaling factor. A significant difference in population scaling factor means was detected across either side of the mean YAP/TAZ ratio (T-Test, n = 4, 5, P ¡0.0001, each mean calculated from 2000 – 5000 cells depending on cell line). C) Nuclear YAP/TAZ concentration against the average cytoplasmic YAP/TAZ scaling factor. No clear relationship was observed across cell lines.

**S.Figure 10:**
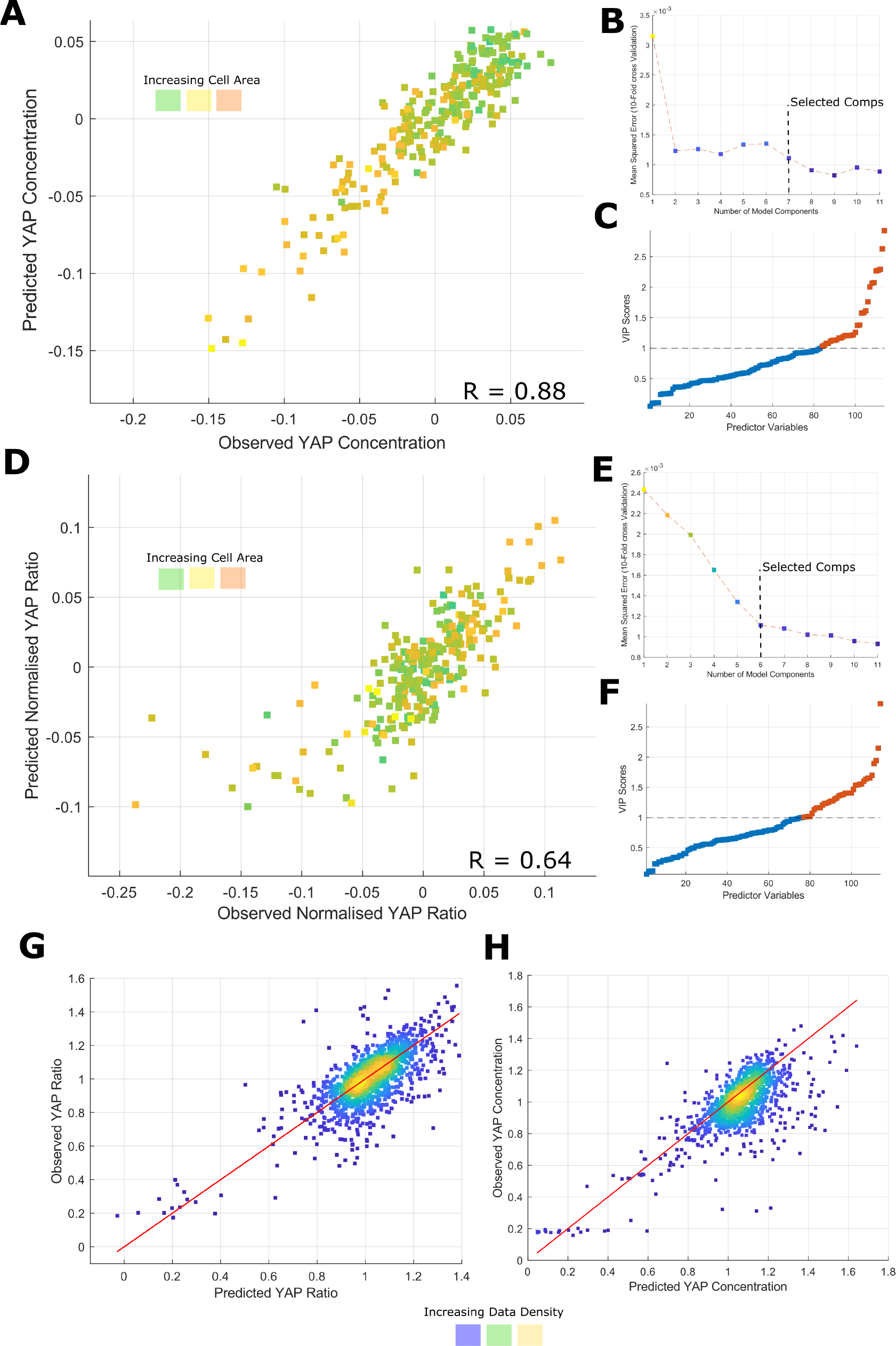
PLSR modelling of YAP/TAZ abundance and ratio in LM2 Cells A) PLSR model predicting well-average YAP/TAZ concentration from morphological and cytoskeletal intensity features in control cells (R = 0.88). The colour reflects the mean cell size in each well. B) Mean squared error evaluated through 10-fold cross validation as a function of component number. The dotted line represents the selected component number. C) Variable importance to projection (VIP) scores for the predictor variables, a score ¿ 1 is considered high and the corresponding feature, important in the prediction; 25% of features significantly contribute to the prediction of YAP/TAZ concentration. D-F) Follows the same pattern as A-C, but relates to the prediction to YAP/TAZ ratio. G) Application of the model in ‘A’ to the prediction of YAP/TAZ concentration in the knockdown states (R = 0.48). The colour represents the density of the data and the red line traces y = x. H) Application of the model in ‘D’ to the prediction of YAP/TAZ ratio in the knockdown states (R = 0.40). The colour represents the density of the data and the red line traces y = x.

**S.Figure 11:**
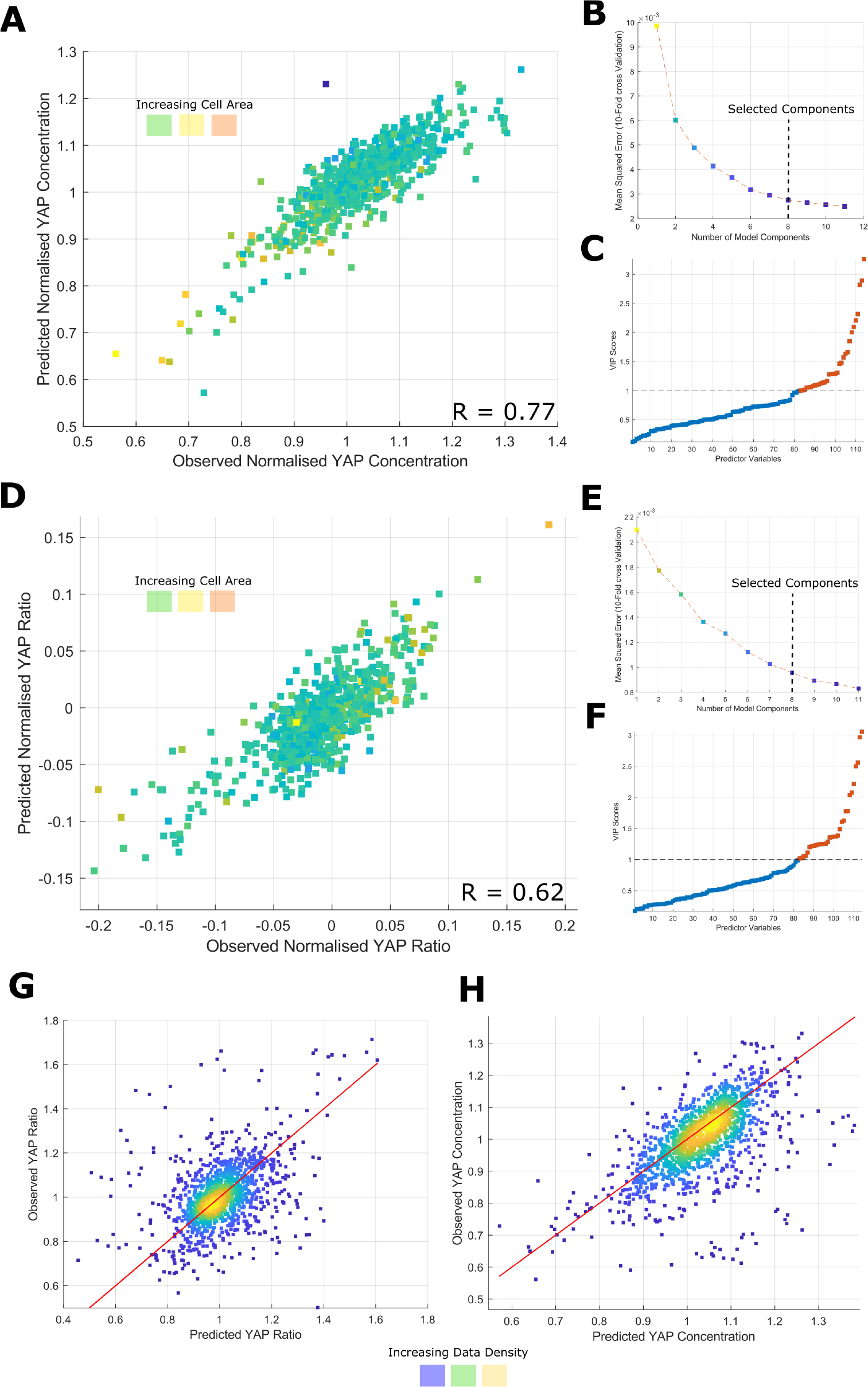
PLSR modelling of YAP/TAZ abundance and ratio in LM2 Cells A) PLSR model predicting well-average YAP/TAZ concentration from morphological and cytoskeletal intensity features in control cells (R = 0.88). The colour reflects the mean cell size in each well. B) Mean squared error evaluated through 10-fold cross validation as a function of component number. The dotted line represents the selected component number. C) Variable importance to projection (VIP) scores for the predictor variables, a score ¿ 1 is considered high and the corresponding feature, important in the prediction; 25% of features significantly contribute to the prediction of YAP/TAZ concentration. D-F) Follows the same pattern as A-C, but relates to the prediction to YAP/TAZ ratio. G) Application of the model in ‘A’ to the prediction of YAP/TAZ concentration in the knockdown states (R = 0.48). The colour represents the density of the data and the red line traces y = x. H) Application of the model in ‘D’ to the prediction of YAP/TAZ ratio in the knockdown states (R = 0.40). The colour represents the density of the data and the red line traces y = x.

